# SuFEx and macrocyclic chelation define orthogonal reactivity domains enabling isotopically versatile radiopharmaceuticals from a single peptide precursor

**DOI:** 10.64898/2026.05.27.728245

**Authors:** Ajay Kumar Sharma, Akhilesh Mishra, Kuldeep Gupta, Sridhar Nimmagadda

**Affiliations:** The Russell H. Morgan Department of Radiology and Radiological Science, Johns Hopkins University School of Medicine, Baltimore, MD, 21287, USA; Department of Oncology, The Sidney Kimmel Comprehensive Cancer Center and the Bloomberg–Kimmel Institute for Cancer Immunotherapy, Baltimore, MD, 21287, USA; Department of Pharmacology and Molecular Sciences, Johns Hopkins University School of Medicine, Baltimore, MD, 21287, USA; Division of Clinical Pharmacology, Department of Medicine, Johns Hopkins University School of Medicine, Baltimore, MD, 21287, USA

## Abstract

The design of radiotheranostic agents has been constrained by a fundamental chemical incompatibility: existing strategies for fluorine-18 incorporation — nucleophilic substitution, silicon-fluoride exchange, aluminum fluoride chelation, and prosthetic conjugation — either require conditions incompatible with macrocyclic metal coordination, consume the chelator site needed for radiometal labeling, or introduce non-native structural appendages that alter pharmacokinetic behavior. Here we show that sulfur(VI) fluoride exchange (SuFEx) chemistry and macrocyclic metal coordination define non-overlapping reactivity domains co-embeddable within a single peptide precursor. An aryl fluorosulfate on tyrosine accepts ¹⁸F under mildly basic conditions; a spatially distinct macrocyclic chelator, DOTA, NOTA, NODAGA, or any compatible macrocycle, independently coordinates diagnostic (^68^Ga, ^64^Cu) or therapeutic (^177^Lu) radiometals under mildly acidic conditions. The labeling pathways proceed independently under mutually compatible conditions, and both preserve receptor-binding affinity. Validated across PD-L1- and CD38-targeting scaffolds, this platform delivers nanomolar target affinity, high radiochemical yields, and matched pharmacokinetics, establishing isotopic orthogonality as a designable, intrinsic property of synthetic molecular radiopharmaceuticals.

## INTRODUCTION

Radiotheranostic agents—molecules that can be used for both disease imaging and targeted radionuclide therapy—have transformed precision oncology by enabling patient stratification, treatment selection and therapeutic monitoring ^1,2^. Clinically established matched radiometal pairs such as [^68^Ga]/[^177^Lu]-DOTA-TATE and [^68^Ga]/[^177^Lu]-PSMA-617 exemplify this power, yet they rely exclusively on radiometal coordination chemistry ^1^. Despite extensive advances in organofluorine and coordination chemistry, a general molecular design that integrates the premier positron emission tomography (PET) isotope Fluorine-18(^18^F) with radiometal labelling capability within a single, chemically unified scaffold has remained elusive^3,4^.

The limitation arises from fundamentally different reactivity requirements that govern ^18^F incorporation and radiometal complexation. Nucleophilic ^18^F substitution requires an activated leaving group, strong bases and rigorously anhydrous conditions to maintain fluoride nucleophilicity, whereas electrophilic ^18^F routes similarly demand harsh or highly specialized reagents ^3^. These constraints stand in sharp contrast to the aqueous, mildly acidic buffers required for the rapid formation of kinetically inert macrocyclic chelates such as DOTA, NOTA or NODAGA, where metal-ion solvation and controlled coordination are essential for thermodynamic and kinetic stability ^4^. Moreover, the short 110-min half-life of ^18^F imposes a strict requirement for late-stage, functionally tolerant chemistry, a persistent challenge in peptide radiolabeling, where densely functionalized, acid–base-sensitive scaffolds offer few intrinsically addressable sites.

Traditional SNAr-based strategies require strongly activated arenes and strongly basic conditions ^3^, whereas clinically used peptide tracers frequently rely on bifunctional prosthetic groups that introduce substantial non-native bulk known to affect lipophilicity, metabolism and in-vivo pharmacokinetics even when receptor affinity is preserved. Aluminium [^18^F]fluoride (AlF) chelation achieves ^18^F incorporation under mild aqueous conditions using NOTA-based chelators, but necessarily occupies the chelator site for fluorine labeling, precluding radiometal coordination from the same scaffold^5^. Aryltrifluoroborate (BF_3_) isotope exchange similarly operates under mild aqueous conditions but requires acidic pH (∼2-4) that overlaps with macrocyclic chelation window and is susceptible to in vivo hydrolysis^6^. These divergent and restrictive labelling pathways have forced radiotheranostic development into parallel synthetic routes for diagnostic and therapeutic agents that differ not only in isotope but also in charge, hydrophilicity and three-dimensional topology, complicating pharmacokinetic harmonization, dosimetry and regulatory approval.

The practical consequences of this incompatibility are exemplified in prostate-specific membrane antigen (PSMA) theranostics. Although ^68^Ga-labelled analogues are available ^7,8^, the ^18^F-labelled tracer ^18^F-DCFPyL has achieved broad clinical adoption in the United States owing to the superior image quality, higher molar activity and reliable biodistribution afforded by cyclotron-produced ^18^F ^9,10^. This clinical divergence highlights how the distinct radiochemical requirements of ^18^F and radiometals, established above, necessarily yield diagnostic and therapeutic agents that differ in charge, hydrophilicity and three-dimensional topology. These structural discrepancies manifest as altered pharmacokinetics, biodistribution, dosimetry and regulatory complexity, reinforcing the need for a unified, orthogonal radiochemistry platform capable of supporting both ^18^F and radiometal labelling from a single precursor without introducing such divergence.

Sulfur(VI) fluoride exchange (SuFEx) chemistry offers an elegant solution. SuFEx exploits selective nucleophilic substitution at sulfur(VI) centres of aryl-sulfonyl fluorides and aryl-fluorosulfates ^11^. The resulting S–F bond is thermodynamically robust yet kinetically tunable, permitting rapid fluoride exchange under mild basic conditions ^12^, well aligned with the constraints of late-stage ^18^F radiochemistry. Although SuFEx-[^18^F] chemistry has emerged as an attractive approach for small-molecule radiofluorination ^13^, with subsequent applications spanning PSMA-targeted imaging agents^14^, fibroblast activation protein inhibitors^15^, and fluorosulfate-modified amino acid tracers^16^, prior implementations were confined to small-molecule substrates and did not address compatibility with macrocyclic peptide architectures, radiometal chelation, or the construction of unified precursors for matched diagnostic and therapeutic isotopes. The ability to integrate SuFEx radiofluorination and radiometal labeling within the same peptide scaffold therefore represents a conceptual advance beyond existing SuFEx radiochemistry. In contrast to silicon-fluoride isotope-exchange strategies (e.g., SiFA-based radiohybrid peptides ^17^) that require prosthetic silicon appendages and exhibit sensitivity to acidic and oxidative environments, SuFEx provides a silicon-free, covalent, and intrinsically chemically orthogonal route to ^18^F incorporation that is mechanistically independent of macrocyclic metal coordination.

We define isotopic orthogonality operationally: two labeling pathways are orthogonal when each proceeds efficiently under its own conditions without compromising the reactive site required by the other, as evidenced by radiochemical yields, product integrity, and preserved biological function. Here we show that SuFEx chemistry can be integrated with macrocyclic chelation to create a chemically orthogonal platform for isotopic versatility in peptide scaffolds. By embedding a tyrosine-derived aryl fluorosulfate as a site for ^18^F incorporation and positioning a macrocyclic chelator on a spatially distinct residue, we generate peptides that can be labelled independently with ^18^F or with radiometals under mutually compatible, mild conditions. The same precursor therefore serves as the source of both diagnostic and therapeutic forms, although the two labelling modes are performed separately. The labeling pathways proceed independently under mutually compatible conditions, each operating within its own chemical window without compromising the functional integrity of the other labeling site, uniting orthogonal fluorine and coordination chemistry within one molecular design. Importantly, this design principle is general (Figure 1). The SuFEx–chelation platform supported efficient radiolabelling when paired with diverse chelators (DOTA, NOTA, NODAGA), accommodated multiple diagnostic and therapeutic isotopes (^68^Ga, ^64^Cu, ^177^Lu) and operated across distinct peptide scaffolds targeting Programmed death ligand 1 (PD-L1) and cluster of differentiation 38 (CD38). This breadth demonstrates that isotopic orthogonality arises from the chemistry itself rather than the particularities of any one ligand, and establishes SuFEx–macrocycle integration as a modular strategy for constructing isotopically versatile radiopharmaceuticals.

**Figure 1.**
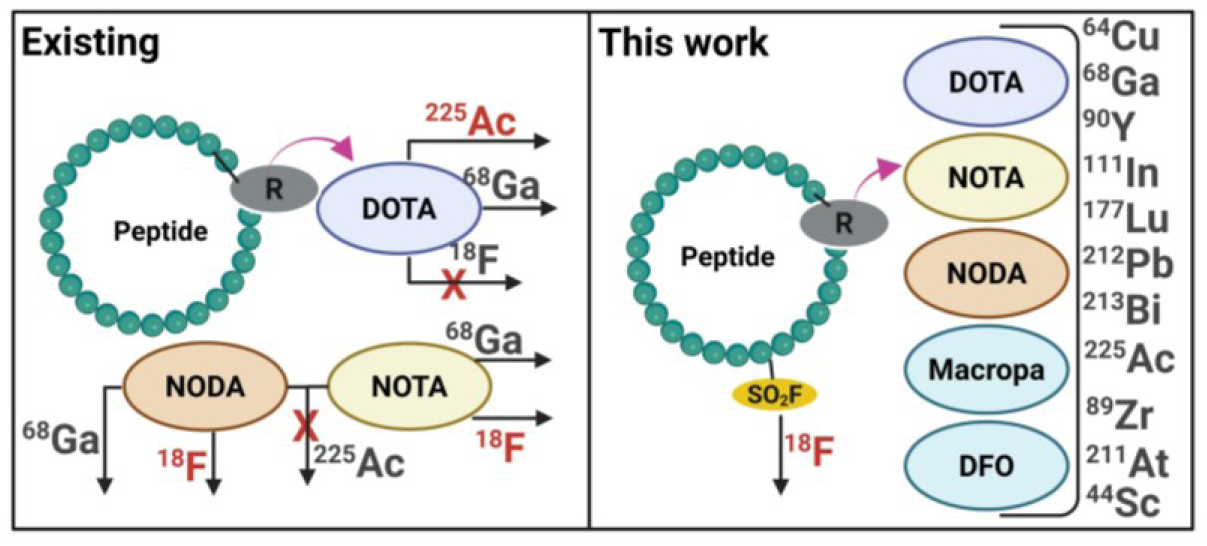
Schematic illustration of the conventional and integrated radiolabeling approaches. The left panel represents the existing peptide-based radiopharmaceutical strategy, where separate chelators are required for radiometal labeling and ^18^F-labeling, limiting the integration of both modalities within a single molecule. The right panel depicts the concept introduced in this work, which enables the incorporation of both radiometal and radiofluorination within the same molecular framework. This integrated design offers a versatile platform for developing dual-modality theranostic agents with potential advantages in personalized nuclear medicine. The figure was created in BioRender. Nimaagadda, S. (2025). https://BioRender.com/j38y259.

We first validated this concept using PD-L1-targeted peptides as an exemplar system. SuFEx-mediated radiofluorination yielded the corresponding ^18^F-labelled tracer with moderate yields under ambient-temperature conditions. In parallel, chelator-based radiometal labeling using DOTA, NOTA and NODAGA facilitated efficient coordination of the positron-emitting radionuclides, ^68^Ga and ^64^Cu, affording the corresponding radiometal analogues in high radiochemical yields. Furthermore, DOTA-based labeling with a therapeutic radionuclide, ^177^Lu, produced the corresponding radiotherapeutic construct in good radiochemical yield. Importantly, the ^18^F-labeling and radiometal-labeling strategies represents two independent, reproducible and mutually compatible pathways. The versatility of this approach was further demonstrated using a CD38-targeting peptide, which showed efficient and compatible radiolabeling with both ^18^F and ^68^Ga. Across all systems, the resulting tracers retained nanomolar binding affinity and exhibited favorable in-vivo pharmacokinetics, indicating that incorporation of both the fluorosulfate moiety and metal chelators is well tolerated within biologically active peptides.

This work extends SuFEx-[¹⁸F] chemistry from small-molecule substrates into the more demanding context of macrocyclic peptide architectures and integrates it with macrocyclic radiometal coordination within a single chemically coherent design. The persistence and function of the fluorosulfate motif within densely functionalized, chelator-bearing macrocyclic peptides reveals a previously undemonstrated dimension of S–F chemistry in complex biomolecular contexts. By integrating radiochemical strategies that have evolved in parallel — organofluorine chemistry and coordination chemistry — this platform provides a generalizable blueprint for multi-isotope radiopharmaceutical development and advances harmonized imaging-to-therapy strategies.

## RESULTS AND DISCUSSION

### Design, synthesis and biochemical characterization of a fluorosulfate-bearing PD-L1 peptide (AJ603)

Macrocyclic peptides that engage PD-L1 with high affinity offer a versatile framework for molecular imaging and targeted radionuclide therapy ^18–20^. Among these, DK221 binds PD-L1 with nanomolar potency through a rigid β-turn motif that presents an exposed tyrosine residue, an attractive locus for late-stage chemical modification^18^. We reasoned that installing a sulfur(VI)–fluoride handle on this phenolic site could create a point of entry for sulfur–fluoride exchange (SuFEx) radiochemistry while preserving the recognition interface.

To realize this design, Fmoc-Tyr(SO₂F)-OH was synthesized (Figures S1–S2) and incorporated into the DK221 sequence by standard solid-phase peptide synthesis, followed by head-to-tail cyclization (Figure 2A, S3). The resulting analogue, AJ603, retains the native sequence but introduces two orthogonal functional groups: an aryl fluorosulfate on tyrosine for ^18^F exchange and an N-allyloxycarbonyl (Alloc) group on lysine for subsequent conjugation. Surface-plasmon-resonance analysis confirmed that AJ603 binds PD-L1 with K_D_ = 46.9 ± 9.7 nM (n = 3; Figure 2B), only modestly weaker than DK221.^18^ This small reduction indicates that the fluorosulfate is electronically benign to the binding interface, validating tyrosine as a tolerated site for SuFEx modification. The Alloc group, however, increases lipophilicity, a feature later optimized through structural refinement.

**Figure 2.**
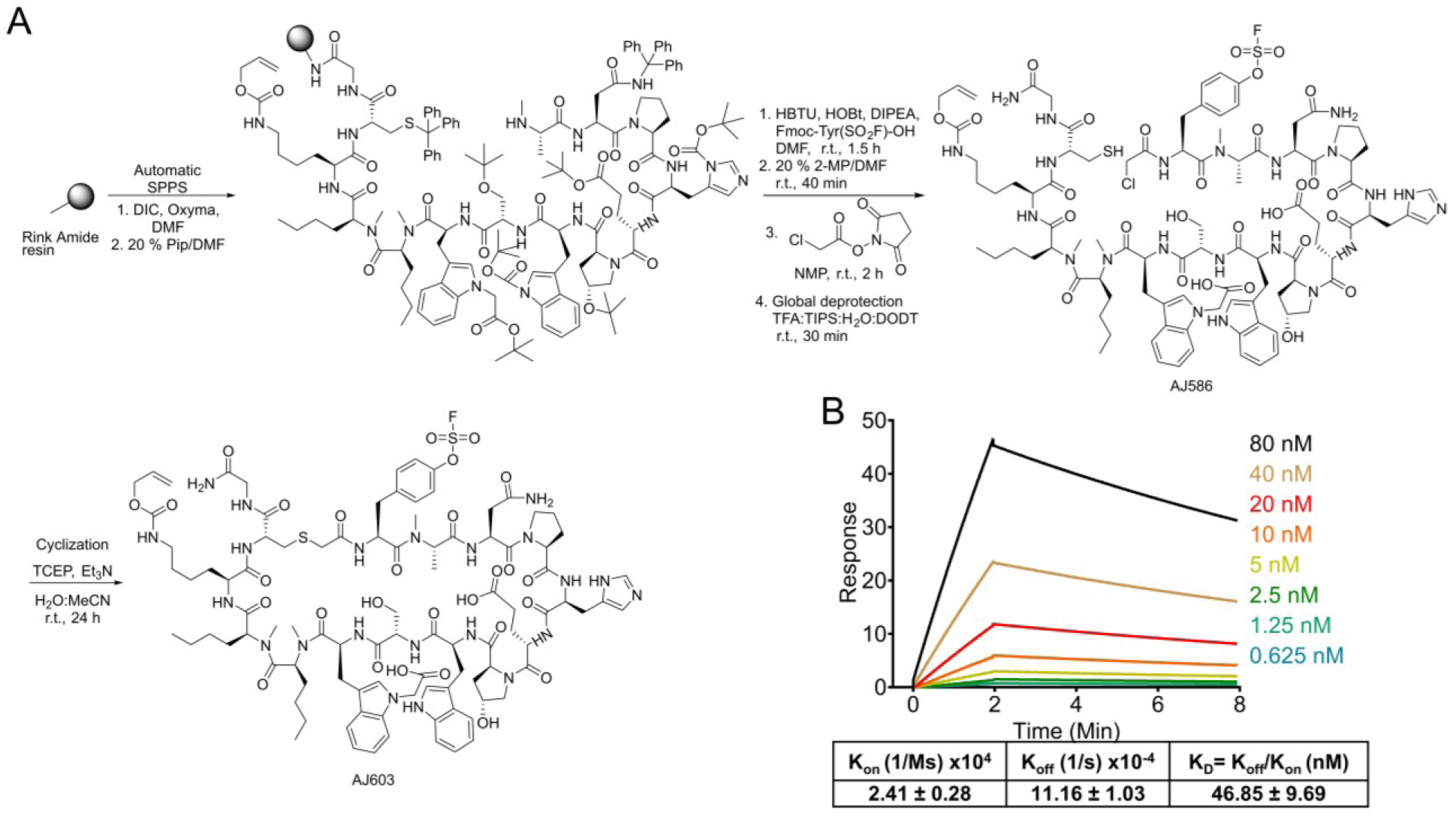
Synthesis of Tyr(SO_2_F) containing PD-L1 binding peptide AJ603 and SPR analysis. **A)** Automatic solid phase peptide synthesis by adding Fmoc-protected amino acids to Rink amide resin using microwave assisted coupling reaction. The created peptidyl resin was treated with cleavage cocktail to obtain deprotected linear peptide. Liner peptide was cyclized in the presence of Et_3_N in water:acetonitrile mixture to obtain AJ603. **B)** Surface plasmon resonance (SPR) analysis showing nanomolar affinity of AJ603 for recombinant PD-L1 protein.

### SuFEx radiofluorination of AJ603 and in-vitro validation

Aryl fluorosulfates undergo fluoride exchange rapidly and selectively under mild basic conditions, a reactivity ideally suited to labile peptide frameworks ^13^. Applying these principles, SuFEx-mediated radiofluorination of AJ603 with [^18^F]fluoride in the presence of K₂CO₃ and Kryptofix 2.2.2 (Figure 3A) furnished [^18^F]AJ603 in a decay-corrected radiochemical yield of 35 ± 16 % (n = 3) and > 98 % radiochemical purity (Figure 3B). Analytical HPLC confirmed co-elution of labelled and non-labelled AJ603 at 14.9 min (Figure 3C). While small-molecule SuFEx radiofluorinations have achieved yields of 83–100% ^13^, the moderate yield observed here is consistent with the more demanding steric and electronic environment of a macrocyclic peptide substrate, demonstrating that the reaction tolerates the polar peptide environment without desulfonylation. The outcome confirms that SuFEx chemistry can extend beyond simple aryl systems to macromolecular substrates while retaining its hallmark operational simplicity.

**Figure 3.**
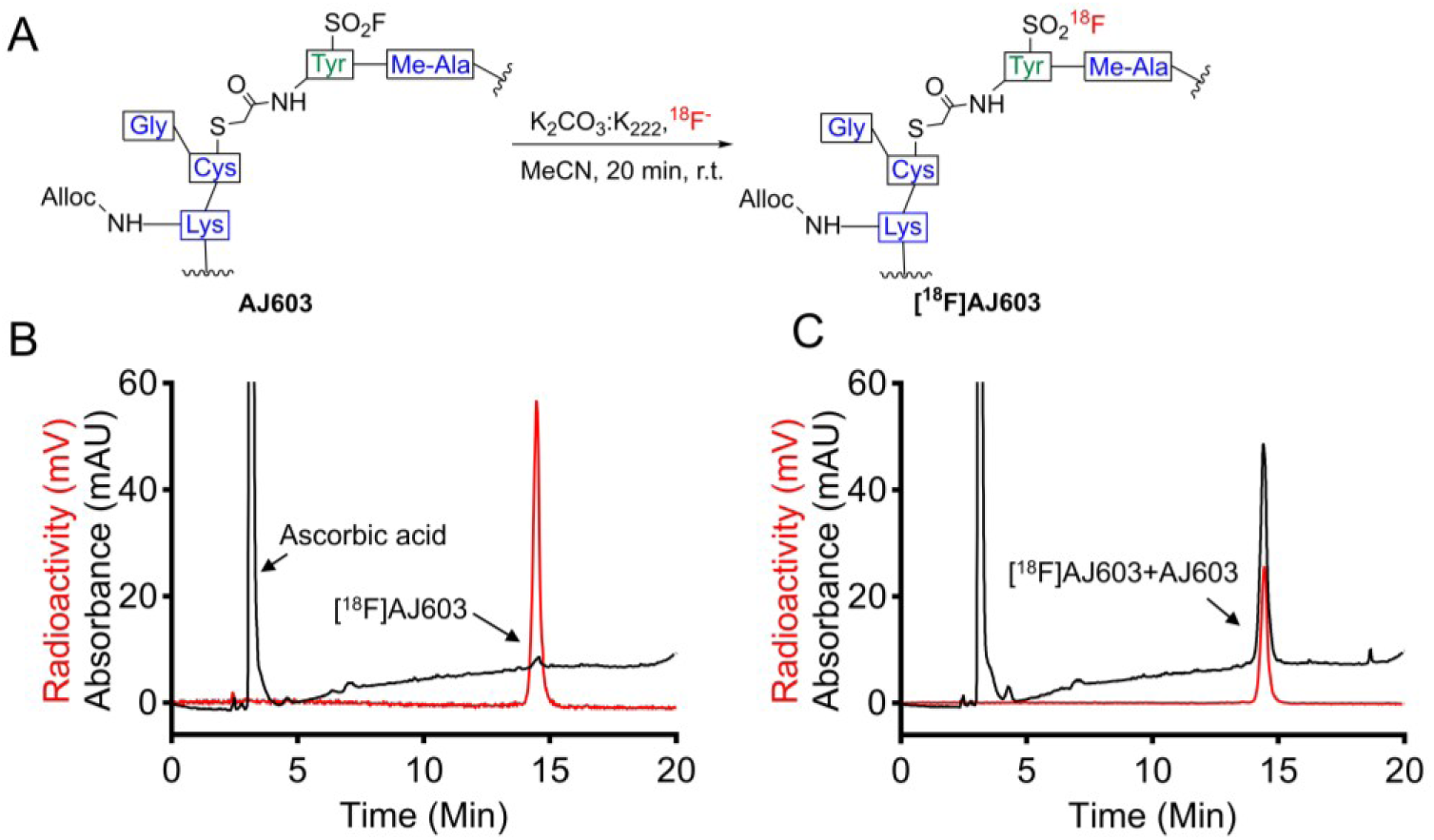
Synthesis and characterization of [^18^F]AJ603. **A)** Radiofluorination scheme of AJ603 employing ^18^F anion, Kryptofix_222_ as phase transfer catalyst in the presence of K_2_CO_3_ as a base and acetonitrile as solvent**. B)** HPLC chromatograms of quality control analysis of formulated radiotracer before conducting studies. **C)** HPLC chromatograms confirming the chemical identity of [^18^F]AJ603 by comparing it with non-radioactive AJ603. The matching retention times and peak profiles between [^18^F]AJ603 and non-radioactive AJ603 confirm the successful and specific incorporation of ^18^F into the AJ603 peptide.

To determine whether the fluorosulfate modification influences PD-L1 recognition, we compared [^18^F]AJ603 uptake in bladder-cancer lines with differential PD-L1 expression. Flow cytometry verified PD-L1^high^ BFTC909 and PD-L1^low^ SCaBER phenotypes (Figure 4A). After incubation with [^18^F]AJ603, BFTC909 cells accumulated 6.53 ± 0.37 % IA versus 0.41 ± 0.08 % IA in SCaBER (Figure 4B), corresponding to a 16-fold selectivity. Competition with 1 µM non-radioactive AJ603 reduced uptake to baseline, confirming receptor-mediated binding (Figure 4B). The data show that aryl-fluorosulfate substitution, despite introducing an electron-withdrawing S(VI) center, leaves the peptide’s recognition properties intact—an essential prerequisite for orthogonal labeling from a single precursor.

**Figure 4.**
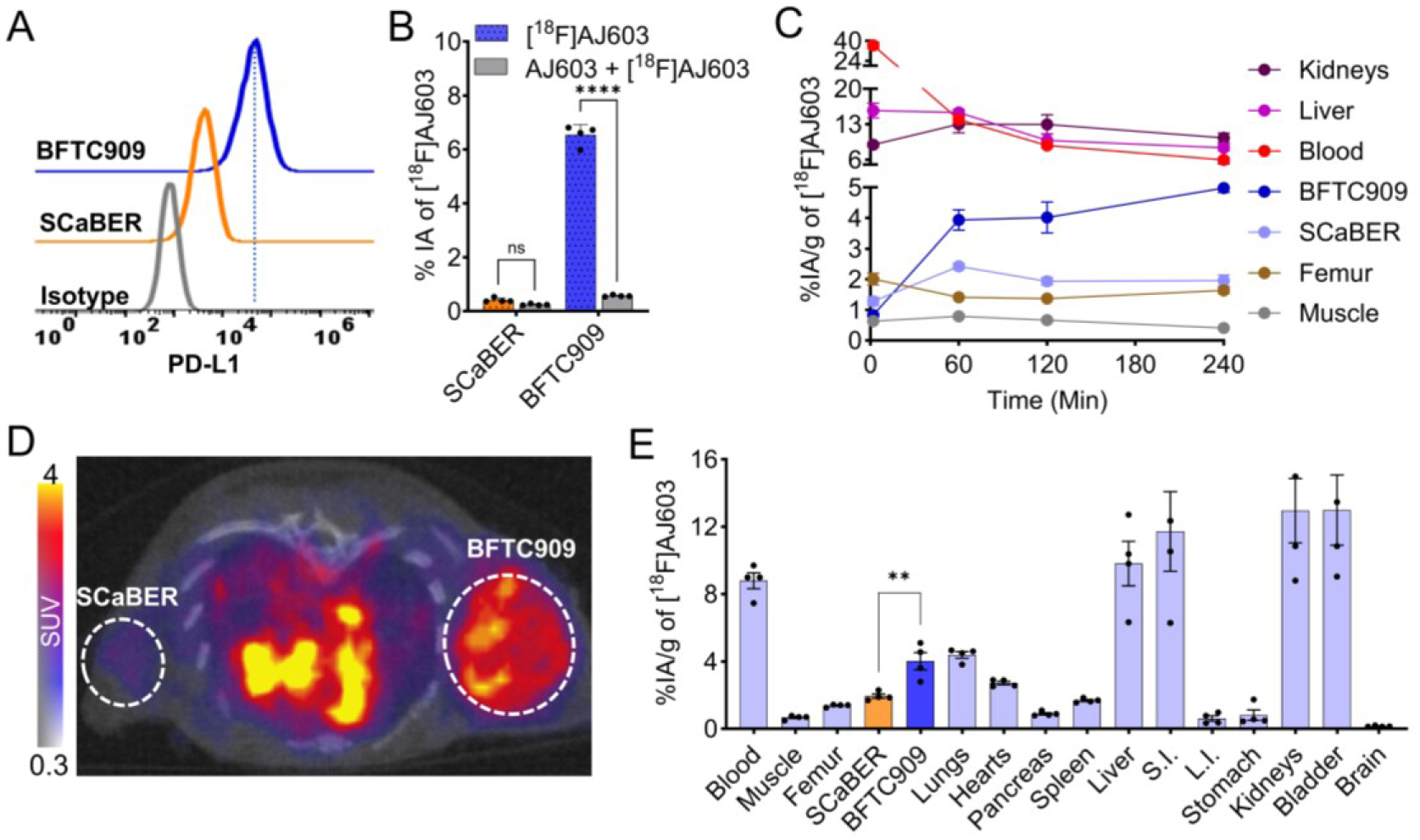
*In vitro* and in vivo assessment of [^18^F]AJ603 in bladder cancer cells and xenograft. **A)** Flow cytometry assay to evaluate cell surface PD-L1 expression in bladder cancer cell lines **B)**[^18^F]AJ603 binding (percent incubated activity, %IA) to bladder cancer cells. Cells were incubated with 1 µCi [^18^F]AJ603 at 4°C for 1 hour. [^18^F]AJ603 uptake is PDL-1 expression dependent, and co-incubation with 1 μM of non-radioactive AJ603 (blocking dose) significantly reduced radiotracer uptake confirming PDL-1 specificity. **C)** Time dependent ex vivo biodistribution data showing accumulation of [^18^F]AJ603 in selected organs of mice bearing bladder cancer xenografts. Full data is in table. **D)** Transaxial section of a PET image acquired with [^18^F]AJ603. PET imaging was performed 120 minutes after intravenous injection of 9.25 MBq (∼250 μCi) of [^18^F]AJ603 in mice bearing bladder cancer xenografts. Tumors are indicated with white circles: ScaBER (upper left flank) and BFTC909 (upper right flank). **E)** Ex vivo biodistribution analysis in tumor-bearing mice at 120 minutes post-injection of [^18^F]AJ603, where S.I. is small intestine and L.I. is large intestine. Data in panels **B** is presented as mean±SD (n=3-4) and in panels **E** is presented as mean±SEM (n=4). Statistical significance was calculated using two-way ANOVA in panel **B** and unpaired *t* test in **E**; ns, *P* ≥ 0.05; **, *P* ≤ 0.01; ****, *P* ≤ 0.0001.

### In-vivo performance and pharmacokinetic behavior of [¹⁸F]AJ603

To assess in-vivo behavior, [^18^F]AJ603 was evaluated in mice bearing PD-L1^high^ (BFTC909) and PD-L1^low^ (SCaBER) xenografts. The goal was to verify PD-L1-dependent accumulation and to identify any pharmacokinetic effects arising from the Alloc and fluorosulfate substituents. Ex-vivo biodistribution revealed early and sustained tumor retention in BFTC909 xenografts, with 3.93 ± 0.34, 4.01 ± 0.51 and 4.98 ± 0.14 %IA g⁻¹ at 60, 120 and 240 min, respectively (Figure 4C; Table S1). Uptake in SCaBER remained significantly lower (2.42 ± 0.10, 1.94 ± 0.13 and 1.96 ± 0.18 %IA g⁻¹), consistent with PD-L1-specific accumulation. The sustained retention implies a receptor-bound residence time compatible with PET imaging windows and confirms that SuFEx modification does not destabilize in-vivo binding.

Background activity in muscle and bone was low, giving tumor-to-muscle ratios of 6.1 ± 0.8 and 12.2 ± 0.3 at 120 and 240 min, respectively (Figure S4). Low femoral activity also indicates minimal defluorination, underscoring the metabolic robustness of the aryl-fluorosulfate bond. PET imaging at 120 min corroborated these findings, with clear delineation of PD-L1^high^ tumors and negligible background (Figure 4D; Figure S5). Elevated uptake in liver (9.8 ± 1.3 %IA g⁻¹) and blood (8.8 ± 0.5 %IA g⁻¹) suggested increased lipophilicity and plasma protein binding, narrowing the optimal imaging window. These data motivated incorporation of a hydrophilic moiety to improve clearance while maintaining orthogonal labelling chemistry.

### Engineering a dual-labelable scaffold through DOTA conjugation (AJ662)

To improve hydrophilicity and enable radiometal incorporation, the Alloc group of AJ603 was removed under palladium catalysis to give AJ661, and the liberated lysine amine was coupled to DOTA to yield AJ662 (Figure 5A; Figure S6). This modification was designed to (i) increase water solubility and (ii) provide a chelating site for radiometal coordination under conditions orthogonal to SuFEx. SPR analysis showed that AJ662 binds PD-L1 with K_D_ = 20.6 ± 13.8 nM (Figure 5B), comparable to AJ603 within experimental error, confirming that DOTA conjugation does not perturb the PD-L1-binding interface.

**Figure 5.**
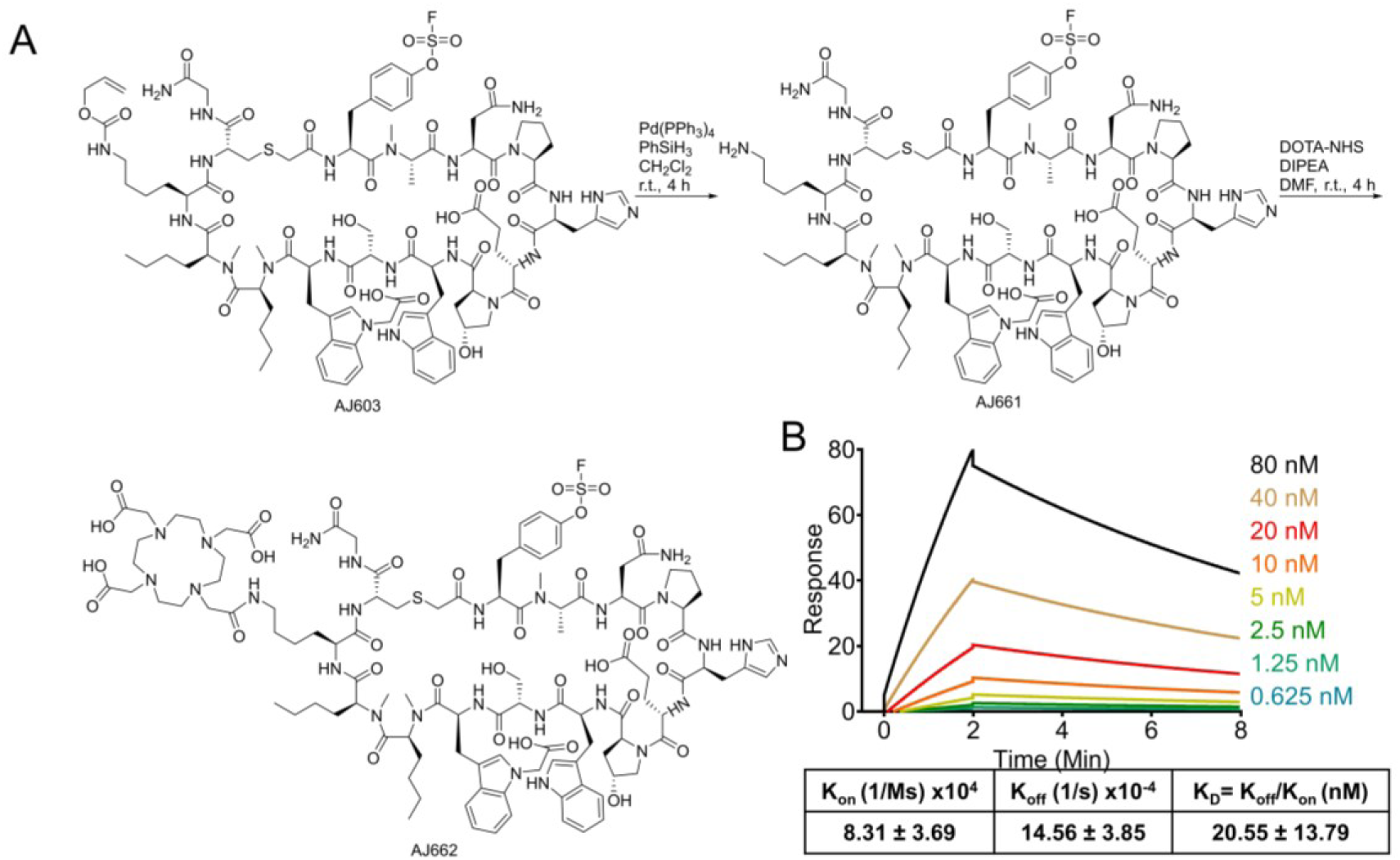
Synthesis of Tyr(SO_2_F) containing PD-L1 binding peptide AJ662 and SPR analysis. **A)** AJ603 was reacted in de-alloc condition, to form AJ661. AJ661 was conjugated with DOTA chelator to obtain AJ662 peptide. **B)** Surface plasmon resonance (SPR) analysis showing nanomolar affinity of AJ662 for recombinant PD-L1 protein.

Mechanistically, the macrocyclic chelator and the aryl fluorosulfate occupy distinct chemical domains: DOTA coordinates radiometals in mildly acidic buffers (pH ≈ 4–5), whereas SuFEx exchange operates under basic conditions. This separation of reactivity implies that both isotopic labelling pathways can proceed independently without mutual interference—a key criterion for achieving isotopic orthogonality within one scaffold.

### Orthogonal radiolabelling of AJ662 with radiometals and fluorine-18

To validate orthogonality, AJ662 was independently labelled via the two chemical routes (Figure 6A). Chelation with ^68^GaCl₃ in acetate buffer (pH 4.5) produced [^68^Ga]AJ662 in > 90 % decay-corrected yield and > 98 % purity with molar specific activity of 16.33 ± 8.00 GBq/µmol (Figure 6B). Co-injection with [^nat^Ga]AJ662 showed identical retention times (Figure 6C), confirming identity. That the ⁶⁸Ga-labelled product co-elutes precisely with its cold counterpart confirms that the fluorosulfate handle survives Ga-68 chelation conditions (pH 4.5, 95°C) chemically intact, providing direct chromatographic evidence of operational orthogonality in the radiometal labeling direction. Radiofluorination using the same SuFEx protocol as AJ603 yielded [^18^F]AJ662 in 42.8 ± 18.6 % yield (n = 5) and > 98 % purity with molar specific activity of 8.35 ± 5.20 GBq/µmol (Figure 6D). The molar activity values reflect the first-generation nature of this platform and are expected to improve with automated synthesis optimization. Co-injection with non-radiolabelled AJ662 confirmed chemical identity (Figure 6E). Comparable yields for AJ662 and AJ603 demonstrate that the presence of a macrocyclic chelator neither perturbs the SuFEx reaction nor competes for fluoride, confirming orthogonality in the radiofluorination direction and completing the bidirectional evidence for the platform’s chemical design principle. Importantly, these reactions are sequential, not simultaneous: the same precursor can generate either isotope variant as needed, but never both in one molecule. This operational independence defines isotopic orthogonality in practice.

**Figure 6.**
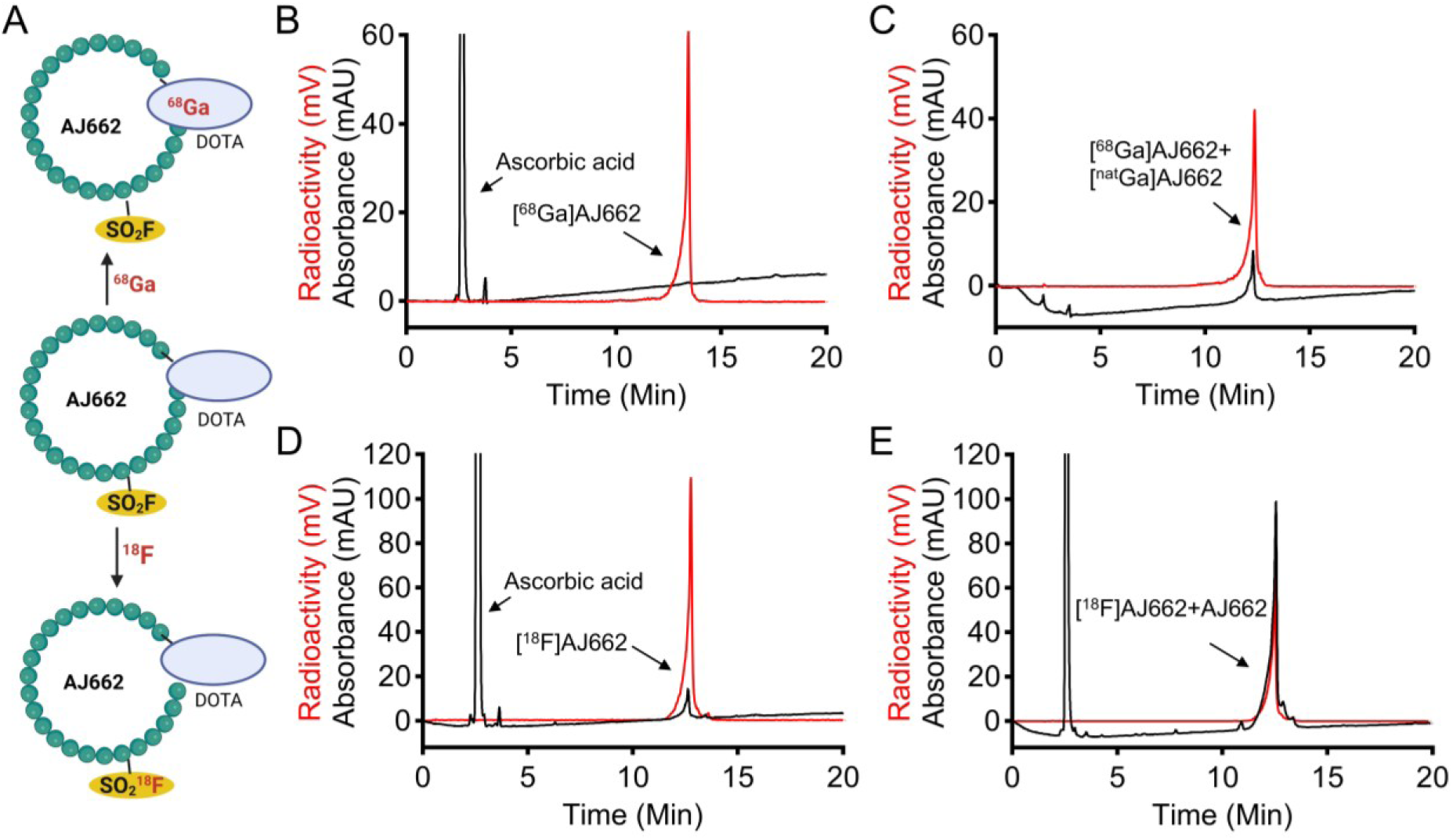
Dual radioisotope labeling of AJ662. **A)** Schematic representation of dual radioisotope labeling of AJ662. Radiometal labeling with Ga-68 was performed on the bifunctional chelator DOTA at the N-terminal of the peptide sequence. Additionally, radiofluorination was facilitated by the fluorosulfate group present on the tyrosine residue in AJ662. **B)** HPLC chromatogram showing the quality control analysis of [^68^Ga]AJ662 before conducting further studies. **C)** HPLC chromatogram confirming the chemical identity of [^68^Ga]AJ662 by comparison with the non-radioactive counterpart [^nat^Ga]AJ662. The matching retention times and peak profiles validate the successful and specific incorporation of Ga-68 into the AJ662 peptide. **D)** HPLC chromatogram showing the quality control analysis of formulated [^18^F]AJ662 before conducting further studies. **E)** HPLC chromatogram confirming the chemical identity of [^18^F]AJ662 by comparison with non-radioactive AJ662. The matching retention times and peak profiles confirm the successful and specific incorporation of F-18 into the AJ662 peptide.

### In-vitro and in-vivo evaluation of [^68^Ga]AJ662

Having confirmed radiochemical compatibility, we next assessed biological performance of [^68^Ga]AJ662. In vitro, the tracer exhibited PD-L1-dependent uptake: 34.6 ± 1.7 % IA in BFTC909 versus 0.77 ± 0.10 % IA in SCaBER (Figure 7B), a ∼45-fold difference and an improvement over [^18^F]AJ603 (∼16-fold). Competition with excess AJ662 reduced uptake to 0.21 ± 0.03 % IA (BFTC909) and 0.06 ± 0.01 % IA (SCaBER), confirming specificity. In mice, [⁶⁸Ga]AJ662 rapidly accumulated in PD-L1^high^ tumors (6.32 ± 0.39 %IA g⁻¹ at 30 min; 7.65 ± 1.08 %IA g⁻¹ at 120 min) with sustained retention, while PD-L1^low^ tumors showed lower, declining uptake (3.70 ± 0.16 → 1.93 ± 0.25 %IA g⁻¹) (Figure 7C–D; Table S2). Background clearance was rapid (muscle 0.55 ± 0.04 %IA g⁻¹; blood 2.08 ± 0.07 %IA g⁻¹ at 120 min), and liver uptake was modest (2.60 ± 0.11 %IA g⁻¹), both markedly improved over [^18^F]AJ603. Tumor-to-muscle ratios increased from 10.3 ± 0.9 (60 min) to 14.2 ± 2.2 (120 min) (Figure S8A–C). Co-injection with cold AJ662 reduced BFTC909 uptake to 2.68 ± 0.42 %IA g⁻¹ at 120 min, while SCaBER uptake was unchanged (2.01 ± 0.04 %IA g⁻¹) (Figure S9A–D). High renal accumulation (12.0 ± 0.4 → 13.2 ± 1.4 %IA g⁻¹, 60–120 min) reflected predominant kidney clearance, a desirable trait that limits non-target background. These data confirm that radiometal labelling preserves peptide integrity and yields superior imaging contrast compared to the initial ^18^F analogue.

**Figure 7.**
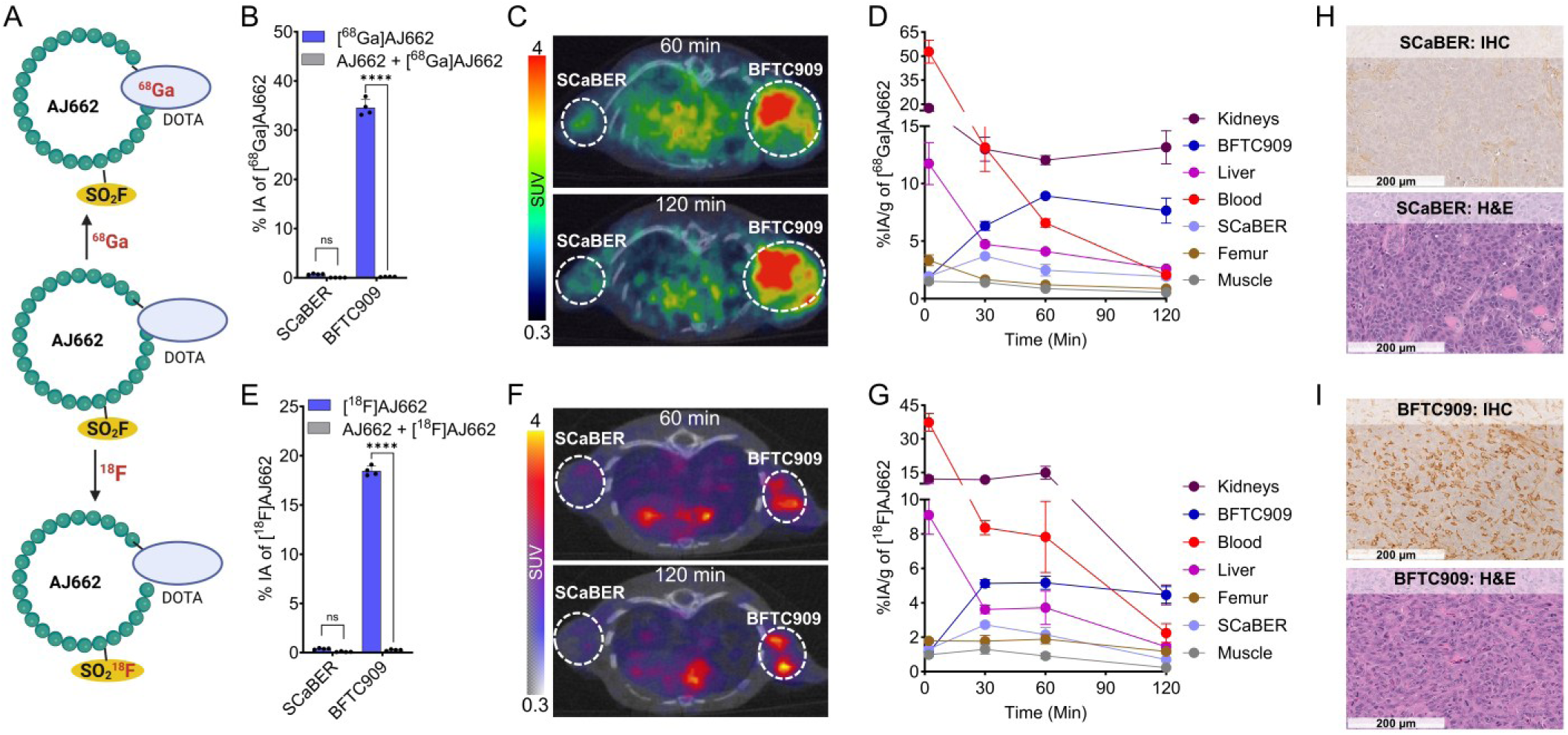
In vitro and in vivo characterization of [^68^Ga]AJ662 and [^18^F]AJ662. **A)** Schematic illustration showing the sites of radiometalation and radiofluorination within the AJ662 molecule. **B)** In vitro uptake of [^68^Ga]AJ662 in bladder cancer cell lines at 4 °C. **C)** In vivo uptake of [^68^Ga]AJ662 in BFTC909 and SCaBER tumor xenograft models evaluated by static PET imaging at 60 and 120 min post-injection. **D)** In vivo time–activity curves of [^68^Ga]AJ662 in BFTC909 and SCaBER tumor xenograft models derived from ex vivo biodistribution data up to 120 min post-injection. **E)** In vitro uptake of [^18^F]AJ662 in bladder cancer cell lines at 4 °C. **F)** In vivo uptake of [^18^F]AJ662 in BFTC909 and SCaBER tumor xenograft models assessed by static PET imaging at 60 and 120 min post-injection. **G)** In vivo time–activity curves of [^18^F]AJ662 in BFTC909 and SCaBER tumor xenograft models derived from ex vivo biodistribution data up to 120 min post-injection. **H, I)** PD-L1 IHC of BFTC909 and SCaBER tumor xenografts confirming target expression. Data in panels **B** and **E** are presented as mean ± SD (n = 4) and data in panels **D** and **G** are presented as mean ± SEM (n = 4). Statistical significance was determined using two-way ANOVA; ns, P ≥ 0.05; ****, P ≤ 0.0001.

### In-vitro and in-vivo evaluation of [^18^F]AJ662

We next examined [¹⁸F]AJ662 to determine whether the DOTA modification affects SuFEx-derived tracer performance. In vitro, uptake in BFTC909 cells was 18.4 ± 0.2 % IA versus 0.39 ± 0.03 % IA in SCaBER, and blocking with excess AJ662 abrogated binding (Figure 7E). In mice, [¹⁸F]AJ662 accumulated selectively in PD-L1^high^ tumors (5.17 ± 0.39 %IA g⁻¹ at 60 min; 4.46 ± 0.48 %IA g⁻¹ at 120 min) with minimal activity in PD-L1^low^ lesions (2.15 ± 0.42 → 0.70 ± 0.08 %IA g⁻¹) (Figure 7F–G; Table S3). Tumor-to-muscle ratios rose from 6.23 ± 1.06 (60 min) to 19.6 ± 2.8 (120 min) (Figure S10). This ratio exceeds that of [^18^F]AJ603 (12.2 ± 0.3), underscoring the positive impact of DOTA on tracer polarity and clearance. Blocking studies confirmed PD-L1 specificity (Figure S11A–D), and immunohistochemistry validated differential PD-L1 expression (Figure 7H–I). Together, these data show that both radiometal and radiofluorine labelling routes yield tracers with excellent target selectivity and complementary imaging performance.

### Comparative pharmacokinetics and optimization insights

Direct comparison of the two isotopic variants revealed similar tumor uptake and clearance kinetics dominated by renal excretion. Notably, hepatic uptake of [^18^F]AJ662 (3.72 ± 0.96 %IA g⁻¹ at 60 min; 1.45 ± 0.15 %IA g⁻¹ at 120 min) was 6–7 fold lower than [^18^F]AJ603 (9.81 ± 1.32 %IA g⁻¹ at 120 min), confirming that removal of Alloc and introduction of DOTA effectively mitigate lipophilicity-driven liver retention. Minor differences in renal activity between [^68^Ga]AJ662 and [^18^F]AJ662 likely reflect coordination chemistry rather than pharmacokinetic disparity. The substantially matched biodistribution profiles of [¹⁸F]AJ662 and [^68^Ga]AJ662 validate the orthogonal labeling strategy, both isotopic forms originate from the same precursor and retain equivalent pharmacokinetic behavior, demonstrating that isotopic identity does not introduce structural divergence.

### Stability and translational robustness of radiolabelled constructs

Radiochemical and metabolic stability are essential for clinical translation. In formulation buffer (saline + 0.1 % Tween-20) at room temperature, both [^68^Ga]AJ662 and [^18^F]AJ662 remained intact for ≥ 2.5 h as assessed by HPLC (Figure 8A–B). In blood samples collected 30 min post-injection and urine up to 120 min, no evidence of demetallation or defluorination was observed. Less than 10 % peptide cleavage occurred in urine by 60 min, slightly higher at 120 min, consistent with normal enzymatic degradation. The combination of in-vitro and in-vivo data confirms that DOTA–Ga coordination and the aryl-fluorosulfate bond are both resistant to hydrolysis under physiological conditions, ensuring that PET signal reflects intact tracer accumulation rather than metabolites. The absence of defluorination in vivo further corroborates the metabolic robustness of the aryl-fluorosulfate bond within the peptide scaffold, complementing the in vitro evidence for orthogonal chemical stability established above. This finding is noteworthy in the context of the broader SuFEx radiofluorination literature, where defluorination leading to bone uptake has been reported as a concern for certain aryl fluorosulfate small-molecule tracers^15^. The favorable stability profile of tyrosine-derived fluorosulfates demonstrated for amino acid tracers^16^ extends in the present work to the substantially more demanding environment of a macrocyclic peptide scaffold bearing a macrocyclic chelator, confirming that the Tyr(SO₂F) motif is metabolically robust across some molecular contexts of increasing complexity.

**Figure 8.**
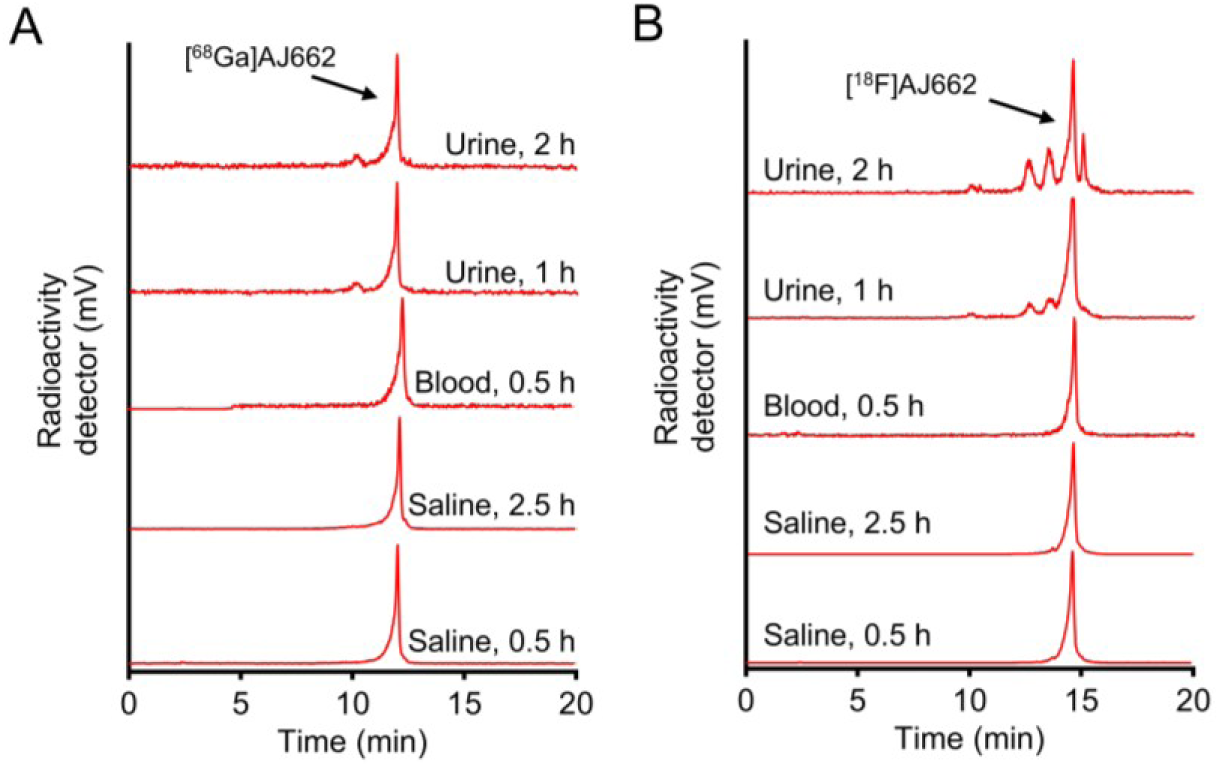
In vitro and in vivo stability assessment of [^68^Ga]AJ662 and [^18^F]AJ662 by HPLC analysis. **A)** In vitro stability of [^68^Ga]AJ662 in saline, with samples collected at 0.5 and 2.5 h, and in vivo stability evaluated using blood samples collected at 0.5 h and urine samples collected at 1 and 2 h post-injection. **B)** In vitro stability of [^18^F]AJ662 in saline, with samples collected at 0.5 and 2.5 h, and in vivo stability evaluated using blood samples collected at 0.5 h and urine samples collected at 1 and 2 h post-injection. All samples were processed and analyzed by radio-HPLC, demonstrating high radiochemical stability of both tracers under in vitro and in vivo conditions.

### Versatile Approach: Multiple Chelator-Modified Derivatives

To demonstrate the versatility of the fluorosulfate-radiometal integrated platform, two additional peptide analogs incorporating alternative chelators, NOTA (AJ698) and NODGA (AJ699), were synthesized (Scheme S1) and fully characterized by MALDI-TOF MS (Figures S12–S13). Both derivatives retained the Tyr(SO₂F) functionality and were efficiently radiolabeled with ^68^Ga, achieving radiochemical yields >90% and radiochemical purities > 98% (Figures S14–S15). In vitro binding studies performed using PD-L1–positive BFTC909 and PD-L1–negative SCaBER cells demonstrated high and specific uptake of both [^68^Ga]AJ698 and [^68^Ga]AJ699 in PD-L1–expressing cells, while uptake was significantly reduced following pre-incubation with excess unlabeled peptide (Figure S16). Importantly, these findings indicate that chelator substitution does not adversely affect radiolabeling efficiency or target binding affinity, underscoring the robustness of the fluorosulfate modification strategy. Collectively, this versatility supports the broad applicability of this platform for the development of fluorosulfate-based radiotracers with flexible chelator selection tailored to specific radiometals and imaging applications.

### Multiple Radiometal Labeling to Enhance Platform Versatility

To further establish the versatility of the fluorosulfate-based peptide platform, AJ662 was radiolabeled with multiple radiometals, including positron emitters for imaging (^68^Ga and ^64^Cu) and a β-emitting radionuclide for radiotherapy (^177^Lu). As discussed earlier, ^68^Ga labeling of AJ662 resulted in high radiochemical yield and excellent in vitro and in vivo performance. Beyond ^68^Ga, AJ662 was successfully radiolabeled with the another positron-emitting radiometal ^64^Cu, achieving high decay-corrected radiochemical yields (>95 %) and high radiochemical purity (Figure S17). In vitro cell-binding assays demonstrated that [^64^Cu]AJ662 exhibited high accumulation in PD-L1–positive BFTC909 cells and minimal uptake in PD-L1–negative SCaBER cells. Importantly, tracer uptake was markedly reduced following pre-incubation with excess unlabeled peptide (Figure S18), confirming PD-L1–specific binding. Similarly, the NOTA- and NODAGA-containing analogs AJ698 and AJ699 were also efficiently radiolabeled with ^64^Cu (Figures S19–S20) and demonstrated high PD-L1 specificity (Figure S21), further supporting the flexibility of the platform with respect to chelator and radiometal selection.

To extend this approach toward therapeutic applications, AJ662 was next radiolabeled with the β-emitting radionuclide ^177^Lu. Radiolabeling proceeded with high efficiency, yielding high radiochemical yield (68 %) and radiochemical purity >95% (Figure S22). In vitro uptake studies in bladder cancer cell lines demonstrated significantly higher accumulation of [^177^Lu]AJ662 in BFTC909 cells compared to SCaBER cells, with PD-L1 specificity further confirmed by competitive blocking using excess unlabeled AJ662 (Figure S23A). In vivo evaluation using ex vivo biodistribution studies in BFTC909- and SCaBER-bearing tumor xenograft models revealed significantly higher tumor uptake of [^177^Lu]AJ662 in PD-L1–positive BFTC909 tumors (6.33 ± 0.86 % IA g^-1^) compared with PD-L1–negative SCaBER tumors (1.85 ± 0.15 %IA g^-1^; n = 3, p = 0.007) at 2 h post-injection. Low uptake was observed in blood and most background tissues, while elevated renal accumulation (11.72 ± 0.64 % IA g^-1^) was consistent with the renal clearance observed with peptide-based radiotracers (Figure S23B).

Collectively, these findings demonstrate that incorporation of the tyrosine-fluorosulfate moiety does not adversely affect radiometal labeling efficiency, in vitro binding, or in vivo target specificity across both diagnostic and therapeutic radionuclides.

### Versatility of the Platform Demonstrated Using an Alternative Target, CD38

To assess whether SuFEx-chelation strategy extends beyond PD-L1-targeted scaffolds, we applied the platform to CD38, a receptor that is highly and uniformly expressed across several hematologic malignancies, including multiple myeloma (MM) and acute myeloid leukemia (AML) ^21^. We previously developed a CD38-targeting peptide derivative, AJ206, and showed its ability to specifically capture CD38-expressing MM cells in vivo ^22^. Because AJ206 contains a tyrosine residue, it provided a direct opportunity to incorporate the tyrosine-fluorosulfate (Tyr–SO₂F) handle without altering the peptide backbone, enabling evaluation of isotopic orthogonality in an unrelated receptor system.

The fluorosulfate-modified construct AJ555 was synthesized to incorporate the Tyr–SO₂F group along with a DOTA chelator and a short PEG_3_ linker (Scheme S2). MALDI-TOF mass spectrometry of the purified peptide confirmed the expected molecular mass (m/z 2272.4, Figure S24) and surface plasmon resonance demonstrated preserved CD38 binding affinity of AJ555 with a dissociation constant (K_D_) of 21.4 nM (Figure S25).

Given the presence of a DOTA chelator, AJ555 was efficiently radiolabeled with ^68^Ga, achieving high decay-corrected radiochemical yields (>90%) and radiochemical purity (>98%) (Figure S26). In vitro uptake studies across multiple myeloma cell lines with varying CD38 expression demonstrated CD38 dependent accumulation of [^68^Ga]AJ555, with highest uptake in MOLP8 cells, followed by RPMI^WT^ cells, and minimal uptake in RPMI^KO^ (CD38 knockout) and U266 cells (Figure S27A). Competitive blocking with excess non-radiolabeled AJ555 resulted in a significant reduction in tracer uptake, and flow cytometry independently confirmed correlation between CD38 expression and radiotracer uptake(Figure S27B).

Next, the in vivo performance of [^68^Ga]AJ555 was evaluated using PET imaging in NSG mice bearing MOLP8 tumor xenografts. PET images revealed robust tumor uptake within 60–120 minutes post-injection, accompanied by rapid clearance from non-target tissues (Figure S27C). Ex vivo biodistribution studies revealed tumor accumulation of 3.94 ± 0.21 % IA g^-1^ at 60 minutes post-injection, which decreased slightly to 2.7 ± 0.29 % IA g^-1^ at 120 minutes, while non-target tissues exhibited low uptake (Table 1, Figure S28A). Notably, a high tumor-to-muscle ratio of 13.57 ± 1.73 was observed at 60 minutes, which increased to 28.30 ± 2.41 at 120 min (Figure S28B). High renal uptake was consistent with peptide-based clearance pathways.

**Table 1.** Chemical characterization of all synthesized peptides.

| Peptide | Chemical formula | Molecular Weight | Exact Mass | Observed MALDI-MS mass |
| --- | --- | --- | --- | --- |
| AJ586 | C<sub>96</sub>H<sub>130</sub>ClFN<sub>22</sub>O<sub>28</sub>S<sub>2</sub> | 2158.8 | 2156.9 | [M + H]<sup>+</sup>; 2159.4 |
| AJ603 | C<sub>96</sub>H<sub>129</sub>FN<sub>22</sub>O<sub>28</sub>S<sub>2</sub> | 2122.3 | 2120.9 | [M + H]<sup>+</sup>; 2122.3 |
| AJ661 | C<sub>92</sub>H<sub>125</sub>FN<sub>22</sub>O<sub>26</sub>S<sub>2</sub> | 2038.3 | 2036.9 | [M + H]<sup>+</sup>; 2038.1 |
| AJ662 | C<sub>108</sub>H<sub>151</sub>FN<sub>26</sub>O<sub>33</sub>S<sub>2</sub> | 2424.7 | 2423.0 | [M + H]<sup>+</sup>; 2424.4 |
| AJ698 | C<sub>104</sub>H<sub>144</sub>FN<sub>25</sub>O<sub>31</sub>S<sub>2</sub> | 2323.6 | 2322.0 | [M + H]<sup>+</sup>; 2323.2 |
| AJ699 | C<sub>107</sub>H<sub>148</sub>FN<sub>25</sub>O<sub>33</sub>S<sub>2</sub> | 2395.6 | 2394.0 | [M + H]<sup>+</sup>; 2395.6 |
| [<sup>nat</sup>Ga]AJ662 | C<sub>108</sub>H<sub>149</sub>FGaN<sub>26</sub>O<sub>33</sub>S<sub>2</sub> | 2492.4 | 2489.9 | [M + H]<sup>+</sup>; 2491.0 |
| [<sup>nat</sup>Ga]AJ698 | C<sub>104</sub>H<sub>142</sub>FGaN<sub>25</sub>O<sub>31</sub>S<sub>2</sub> | 2391.3 | 2388.9 | [M + Na]<sup>+</sup>; 2411.6 |
| [<sup>nat</sup>Ga]AJ699 | C<sub>107</sub>H<sub>145</sub>FGaN<sub>25</sub>O<sub>33</sub>S<sub>2</sub> | 2462.3 | 2460.9 | [M + H]<sup>+</sup>; 2462.0 |
| [<sup>nat</sup>Cu]AJ662 | C<sub>108</sub>H<sub>149</sub>CuFN<sub>26</sub>O<sub>33</sub>S<sub>2</sub> | 2486.2 | 2484.0 | [M + Na]<sup>+</sup>; 2486.8 |
| [<sup>nat</sup>Cu]AJ698 | C<sub>104</sub>H<sub>142</sub>CuFN<sub>25</sub>O<sub>31</sub>S<sub>2</sub> | 2385.1 | 2382.9 | [M + Na]<sup>+</sup>; 2403.4 |
| [<sup>nat</sup>Cu]AJ699 | C<sub>107</sub>H<sub>145</sub>CuFN<sub>25</sub>O<sub>33</sub>S<sub>2</sub> | 2456.1 | 2453.9 | [M + Na]<sup>+</sup>; 2475.1 |

To evaluate SuFEX radiofluorination on this scaffold, AJ555 was subsequently subjected to late-stage ^18^F incorporation using K₂CO₃, kryptofix® 222, and acetonitrile. [^18^F]AJ555 was obtained with approximately 30% decay-corrected radiochemical yield and high radiochemical purity (>98%) (Figure S29). In vitro uptake of [^18^F]AJ555 mirrored CD38 expression levels and was suppressed by blocking (Figure S30A), confirming retention of target specificity following fluorination. In vivo PET imaging with [^18^F]AJ555 in MOLP8 tumor–bearing mice revealed clear tumor visualization at 60 minutes post-injection, with low background activity in muscle (Figure S30B). Accumulation in the liver, likely reflecting the lipophilicity of AJ555 analog, along with high renal uptake indicative of rapid clearance.

Collectively, these in vitro and in vivo data establish that incorporation of the Tyr–SO₂F moiety and macrocyclic chelator is well tolerated within a CD38-targeting peptide and without functional compromise. Demonstrating isotopic orthogonality in a receptor system distinct from PD-L1 confirms that the SuFEx-chelation pairing is not sequence- or target specific, but rather constitutes a modular chemical strategy adaptable across multiple peptide architectures and oncologic targets.

## Conclusion

AJ662 establishes a dual-functional peptide framework in which an aryl-fluorosulfate and a macrocyclic chelator operate as electronically and spatially orthogonal reactivity domains, enabling independent incorporation of ^18^F-or radiometals from a single precursor. Both labelling pathways proceed efficiently and preserve high target affinity and in vivo targeting. The improved tumor-to-background ratios observed for AJ662 relative to AJ603 illustrate that isotopic orthogonality can be achieved while permitting deliberate pharmacokinetic tuning within a unified scaffold.

More broadly, this work extends SuFEx-[^18^F] chemistry from small-molecule substrates to complex peptide architectures and integrates it with macrocyclic radiometal coordination in a chemically coherent design. The compact aryl-fluorosulfate motif enables a mild, site-specific late-stage ^18^F incorporation, while the macrocyclic chelator domain accommodates diagnostic and therapeutic radiometals within the same molecular framework. By systematically integrating radiochemical manifolds that have largely evolved in parallel, this platform provides a generalizable blueprint for multi-isotope radiopharmaceutical design and advancing harmonized imaging-to-therapy strategies.

## EXPERIMENTAL SECTION

### Chemicals

All Fmoc protected amino acids, TIPS, HOBT, and HBTU were obtained from Chem-impex. Rink amide resin and unnatural amino acids, 1,1′-Sulfonyldiimidazole were purchased from Ambeed, while DIPEA, TFA, DODT, DMF and acetonitrile were obtained from Sigma-Aldrich.

### Chemical Synthesis of Fmoc-Tyr(SO_2_F)-OH

Fmoc-Tyr(SO_2_F)-OH was synthesized by reacting SO_2_F_2_ gas (generated in situ by reacting 1,1′-Sulfonyldiimidazole and KF in the presence of TFA) with Fmoc-Tyr-COOH in the presence of triethylamine in DCM for 12 hours at room temperature.^23^ Volatiles were removed under reduced pressure and crude was used in peptide synthesis without any purification. Purity was assessed by RP-HPLC system using a semi-preparative C-18 Luna column (5 mm, 10 x 250 mm Phenomenex, Torrance, CA). The HPLC condition was gradient elution starting with 5% acetonitrile: water (0.1% TFA) and reaching 95% acetonitrile: water (0.1% TFA) in 20 min and continued isocratic mixture of 95% acetonitrile: water (0.1% TFA) till 25 min at a flow rate of 5 mL/min. The elution time (RT) of Fmoc-Tyr(SO_2_F)-OH from HPLC was 18.9 min. The product was characterized by MALDI-TOF-MS. Theoretical Chemical formula: C_24_H_20_FNO_7_S; Exact Mass: 485.1; Molecular Weight: 485.5; Therotical MALDI-MS mass [M + Na]^+^: 508.1; Observed MALDI-MS mass [M + Na]^+^: 508.4.

### Chemical Synthesis of AJ603

AJ603 was chemically synthesized by standard Fmoc solid-phase peptide synthesis on Rink Amide resin employing automatic and manual solid phase peptide synthesis in a 0.1 mmol scale. Automatic coupling was carried out in Liberty Blue peptide synthesizer using DIC (0.5 mmol), Oxyma (0.5 mmol) and Fmoc-AA-OH (0.5 mmol) in DMF at 90 °C under microwave for 2 min. Fmoc group was deprotected using 20% piperidine in DMF at 90 °C under microwave for 1 min. Manual coupling reaction was carried out using HBTU (0.5 mmol), HOBT (0.5 mmol), DIPEA (0.5 mmol) and Fmoc-Tyr(SO_2_F)-OH (0.5 mmol) in DMF at room temperature for 1.5 h. Fmoc group was deprotected using 20% 2-methylpiperidine in DMF for 40 min at room temperature. Chloro-acetyl was conjugated on solid phase by reacting Chloro-acetyl-NHS ester (80 mg) in N-methyl-2-pyrrolidone with free amine of peptidyl resin for 2 hours at room temperature. Once sequence was completed on resin, peptidyl-resin was treated with 4 mL of cleavage cocktail (TFA:TIPS:DODT:H_2_O; 92.5:2.5:2.5:2.5) for 30 min at room temperature. Cleaved reaction mixture was filtered and precipitated with diethyl ether to obtained linear peptide AJ586 as a white solid and characterized by MALDI-TOF mass spectrometry. Theoretical Chemical formula: C_96_H_130_ClFN_22_O_28_S_2_; Exact Mass: 2156.9; Molecular Weight: 2158.8; Therotical MALDI-MS mass [M + H]^+^: 2157.9; Observed MALDI-MS mass [M + H]^+^: 2159.4.

For cyclization,^22^ 50 mL of water:acetonitrile (1:1) was added in round bottomed flask and 50 mg of tris(2-carboxyethyl)phosphine (TCEP), followed by 200 µL of triethylamine was added. Next, 50 mg of AJ586 peptide was dissolved in 4 mL of water:acetonitrile (1:1) and added slowly in reaction mixture over an hour. Reaction mixture was stirred at room temperature for 24 h. Reaction was quenched with TFA, volatiles were removed and purified on a reversed phase high performance liquid chromatography (RP-HPLC) system using a preparative C-18 Phenomenex column (5 mm, 21.5 x 250 mm Phenomenex, Torrance, CA). The HPLC condition was gradient elution started with 10% acetonitrile: water (0.1% TFA) and reached at 60% acetonitrile: water (0.1% TFA) in 30 min followed by next 10 min at 60% acetonitrile: water (0.1% TFA) at a flow rate of 8 mL/min. The product AJ603 was collected at RT ∼33.9 min. Acetonitrile was evaporated under reduced pressure and lyophilized to form an off-white powder with 15 % yield (8 mg). AJ603 was characterized by MALDI-TOF-MS. Theoretical Chemical formula: C_96_H_129_FN_22_O_28_S_2_; Exact Mass: 2120.9; Molecular Weight: 2122.3; Therotical MALDI-MS mass [M + H]^+^: 2121.9; Observed MALDI-MS mass [M + H]^+^: 2122.3.

### Chemical synthesis of AJ662, AJ698 and AJ699

6 mg of AJ603 was dissolved with 6 mg of Pd(PPh_3_)_4_ in anhydrous DCM. PhSiH_3_ (3 molar equivalent) was added in the reaction mixture and stirred for 4 h at room temperature. Volatiles were removed and reaction mixture purified on a reversed phase high performance liquid chromatography (RP-HPLC) system using a preparative C-18 Phenomenex column (5 mm, 21.5 x 250 mm Phenomenex, Torrance, CA). The HPLC condition was gradient elution started with 10% acetonitrile: water (0.1% TFA) and reached at 60% acetonitrile: water (0.1% TFA) in 30 min followed by next 10 min at 60% acetonitrile: water (0.1% TFA) at a flow rate of 8 mL/min. The product AJ661 was collected at RT ∼30.5 min. Acetonitrile was evaporated under reduced pressure and lyophilized to form an off-white powder with 75 % yield.

Next, Intermediate was independently reacted with NHS ester of chelators (AJ662; DOTA-NHS, AJ698; NOTA-NHS and AJ699; NODAGA-NHS) in the presence of DIPEA (10 molar equivalent) in DMF at room temperature for 3 h. Volatiles were removed and reaction mixture purified on a reversed phase high performance liquid chromatography (RP-HPLC) system using a preparative C-18 Phenomenex column (5 mm, 21.5 x 250 mm Phenomenex, Torrance, CA). The HPLC condition was gradient elution started with 10% acetonitrile: water (0.1% TFA) and reached at 60% acetonitrile: water (0.1% TFA) in 30 min followed by next 10 min at 60% acetonitrile: water (0.1% TFA) at a flow rate of 8 mL/min. Acetonitrile was evaporated under reduced pressure and lyophilized to form an off-white powder with around 70-80 % yields. These peptides were characterized by MALDI-TOF-MS and data is in **table 1**.

### Chemical Synthesis of AJ555

AJ555 was chemically synthesized by standard Fmoc solid-phase peptide synthesis on Rink Amide resin employing manual solid phase peptide synthesis in a 0.1 mmol scale ^24^. Coupling reaction was carried out using HBTU (0.5 mmol), HOBT (0.5 mmol), DIPEA (0.5 mmol) and Fmoc-AA-OH (0.5 mmol) in DMF at room temperature for 1.5 h. Fmoc group was deprotected using 20% 2-methylpiperidine in DMF for 40 min at room temperature. Fmoc-PEG_3_-COOH and (^t^Bu)_3_-DOTA-COOH was also incorporated using above mentioned coupling conditions. Once sequence was completed on resin, peptidyl-resin was treated with 4 mL of cleavage cocktail (TFA:TIPS:DODT:H_2_O; 92.5:2.5:2.5:2.5) for 4 h at room temperature. Cleaved reaction mixture was filtered and precipitated with diethyl ether to obtained linear peptide AJ551 as a white solid.

For cyclization^22,25^, 50 mg of AJ551 peptide was dissolved in 50 mL of water:acetonitrile (1:1) and treated with 60 mg of tris(2-carboxyethyl)phosphine (TCEP), followed by 200 µL of Et_3_N. 15 mg of 1,3,5-Tris(bromomethyl)benzene (TBMB) was dissolved in acetonitrile and added slowly in reaction mixture over an hour. Reaction mixture was stirred at room temperature for 24 h. Reaction was quenched with TFA, volatiles were removed and purified on a reversed phase high performance liquid chromatography (RP-HPLC) system using a preparative C-18 Phenomenex column (5 mm, 21.5 x 250 mm Phenomenex, Torrance, CA). The HPLC condition was gradient elution started with 10% acetonitrile: water (0.1% TFA) and reached at 60% acetonitrile: water (0.1% TFA) in 40 min at a flow rate of 8 mL/min. The product AJ555 was collected at RT ∼21.5 min. Acetonitrile was evaporated under reduced pressure and lyophilized to form an off-white powder with 25 % yield. The sequence of AJ555 peptide is DOTA-PEG_3_-ACVPCADFPIWY(SO_2_F)C-NH_2_ with TBMB-based cyclization. This peptide was characterized by MALDI-TOF-MS. Theoretical Chemical formula: C_103_H_143_FN_20_O_29_S_4_; Exact Mass: 2270.92; Molecular Weight: 2272.63; Therotical MALDI-MS mass [M + H]^+^: 2271.9; Observed MALDI-MS mass [M + H]^+^: 2272.8.

### General Method for Affinity Measurements by Surface Plasmon Resonance (SPR)

The affinity of AJ603 and AJ662 for human PD-L1 recombinant protein and the affinity of AJ555 for human CD38 recombinant proteins were evaluated by SPR. The experiments were conducted using a Biacore T200 instrument with a CM5 chip at 25 °C. The ligands used were His-Tagged human PD-L1 (R&D systems, catalog# 9049-B7, 35.5 kDa, 0.5 mg/ml stock concentration) and His-Tagged human CD38 (R&D systems, catalog # 2404-AC, 43 kDa, 0.5 mg/ml stock concentration), which was immobilized onto the CM5 chip. AJ662 (2424.7 Da, 10 mM stock concentration) and AJ603 (2122.4 Da, 10 mM stock concentration) for PD-L1 and AJ555 (2272.6 Da, 10 mM stock concentration) for human CD38 were used as the analyte, which flowed over the ligand immobilized surface. FC2 was used as the experimental flow cell, while FC1 served as the reference. Anti-His antibody (1 mg/ml stock concentration) was immobilized on both FC1 and FC2 using standard amine coupling chemistry. The immobilization running buffer used was PBS-P (20 mM phosphate buffer pH 7.4, 137 mM NaCl, 2.7 mM KCl, 0.05% v/v surfactant P20). Human PD-L1 or human CD38 was captured on FC2 to a level of ∼1200 RU and ∼860 RU, respectively in independent experiments. PBS-P was used as the capture running buffer. Based on these captured response values, theoretical R_max_ values were calculated and are 71.7 for AJ603, 82.0 for AJ662 and 45.5 for AJ555. The R_max_ values assume 1:1 interaction mechanism. Overnight kinetics were performed for all analytes in the presence of PBS-P+5% DMSO. The flow rate of all analyte solutions was maintained at 50 μL/min. The contact and dissociation times used were 120s and 360s, respectively. Surface regeneration was achieved by injecting glycine pH 1.5 for 20 seconds, which takes away all captured ligands onto FC2. Fresh ligands were captured at the beginning of each injection cycle. The analyte concentrations injected ranged from 80 nM down to 0.625 nM for PD-L1 with two-fold serial dilutions and 300 nM down to 1.2 nM for CD38 with three-fold serial dilutions, and all analytes were injected in duplicate.

### General Methods For F-18 Labeling

The one step radiofluorination synthesis was adapted from literature^13^ and modified for our purpose. Briefly, QMA sep-pack was preconditioned by eluting with potassium bicarbonate solution (0.5 M) and water (5 mL) followed by dried using flushing with air. No-carrier-added [^18^F]fluoride was produced in cyclotron. Around 100 mCi of [^18^F]F was trapped onto the preconditioned QMA column and washed with 5 mL water. Activity was eluted with 1 mL mixture of Potassium carbonate (K_2_CO_3_, 5 mg), 2.2.2-cryptand (5 mg) solution. Next, eluted activity was azeotropically dried at 110 °C using acetonitrile. Dried F-18 was redissolved in anhydrous acetonitrile and incubated with 100 µg of peptide with around 40 mCi of radioactivity in total volume of 0.5 mL at room temperature for 25 min. Reaction mixture was diluted with water and ascorbic acid, trapped on C-18 sep-pack column and washed with water. Mixture was eluted with ethanol and added directly in 0.1% tween20 in saline with 100 µg of ascorbic acid and incubated at room temperature for 5 min. Next, mixture was purified on a reversed phase high performance liquid chromatography (RP-HPLC) system using a semi-preparative C-18 Luna column (5 mm, 10 x 250 mm Phenomenex, Torrance, CA). The HPLC condition was gradient elution started with 20% acetonitrile: water (0.1% formic acid) and reached at 60% acetonitrile: water (0.1% formic acid) in 20 min at a flow rate of 5 mL/min. The radiolabeled product [^18^F]AJ603 was collected at RT ∼14.4 min, [^18^F]AJ662 was collected at RT ∼12.6 min and [^18^F]AJ555 was collected at RT ∼13.2 min with decay corrected radiochemical yield of around 30-40%. Desired radiolabeled fraction was concentrated under stream of N_2_ at 60 °C, formulated in 0.1% tween in saline which was used for in vitro and in vivo studies. The whole radiolabeling process for each peptide was done in ∼80-90 min.

### Protocol For Labeling with Cold Ga

Cold Ga labeling of AJ662 was carried out by adding 20 µL of aqueous 0.1M [^nat^Ga]GaCl_3_ solution and 0.6 mL of 0.1 M HCl to a stirred solution of peptides (0.1 mg) in 200 µL of 1M NaOAc buffer (pH 5.0) in a reaction vial. The reaction mixture was incubated for 30 min at 95 °C in a temperature-controlled heating block and purified on a RP-HPLC system using a semi-preparative C-18 Luna column (5 mm, 10 x 250 mm Phenomenex, Torrance, CA). The HPLC conditions were gradient elution starting with 20% acetonitrile: water (0.1% formic acid) and reaching 60% acetonitrile: water (0.1% formic acid) in 20 min at a flow rate of 5 mL/min. The products were collected and acetonitrile was evaporated under reduced pressure and lyophilized to form an off-white powder, which was characterized by MALDI-TOF-MS. Theoretical Chemical formula: C_108_H_149_FGaN_26_O_33_S_2_; Exact Mass: 2489.9; Molecular Weight: 2492.4; Therotical MALDI-MS mass [M + H]^+^: 2490.9; Observed MALDI-MS mass [M + H]^+^: 2491.0.

### General method for labeling with cold Cu

Cold Cu labeling of peptides were carried out by adding 20 µL of aqueous 0.1M [^nat^Cu]CuSO_4_ solution and peptides (0.1 mg) in 200 µL of 0.1 M NaOAc buffer (pH 5.0) in a reaction vial. The reaction mixture was incubated at 70 °C for 60 min in a temperature-controlled heating block and purified on a RP-HPLC system using a semi-preparative C-18 Luna column (5 mm, 10 x 250 mm Phenomenex, Torrance, CA). The HPLC conditions were gradient elution starting with 20% acetonitrile: water (0.1% formic acid) and reaching 60% acetonitrile: water (0.1% formic acid) in 20 min at a flow rate of 5 mL/min. The products were collected and acetonitrile was evaporated under reduced pressure and then were characterized by MALDI-TOF-MS.

### Protocol For Ga-68 Radiolabeling

The ^68^Ge/^68^Ga generator was manually eluted using 6 mL of 0.1M HCl (Ultrapure trace-metal-free) in four different fractions (2.4 mL, 1 mL, 1 mL, and 1.4 mL). To a microcentrifuge vial (1.5 mL) containing 200 μL of 1 M NaOAc buffer (pH = 5) and 40 µg of peptide (around 20 nmoles), 8-10 mCi of [^68^Ga]GaCl_3_ in 0.6 mL from the second fraction was added. The reaction mixture was incubated for 12 min at 95 °C in a temperature-controlled heating block and purified on a RP-HPLC system using a semi-preparative C-18 Luna column (5 mm, 10 x 250 mm Phenomenex, Torrance, CA). The HPLC conditions were gradient elution starting with 20% acetonitrile: water (0.1% formic acid) and reaching 60% acetonitrile: water (0.1% formic acid) in 20 min at a flow rate of 5 mL/min. The radiolabeled product was collected with decay-corrected radiochemical yield of >90 %. The desired radiolabeled fraction was concentrated under a stream of N_2_ at 60 °C, formulated in 0.1% Tween^20^ in saline, and used for *In vitro* and *In vivo* studies. The whole radiolabeling process was completed in approximately 30 min. Quality control was performed on the same HPLC system using the same HPLC gradient as described above.

### General method for Cu-64 radiolabeling

The ^68^Cu was obtained from University of Wisconsin and dissolved in 0.1 M NaOAc (pH 5.0). To a microcentrifuge vial (1.5 mL) containing 200 μL of 0.1 M NaOAc buffer (pH = 5) and 20 µg of Peptide (around 10 nmoles), 1 mCi of [^64^Cu]Cu(OAc)_2_ was added. The reaction mixture was incubated for 60 min at 70 °C in a temperature-controlled heating block and purified on a RP-HPLC system using a semi-preparative C-18 Luna column (5 mm, 10 x 250 mm Phenomenex, Torrance, CA). The HPLC conditions were gradient elution starting with 20% acetonitrile: water (0.1% formic acid) and reaching 60% acetonitrile: water (0.1% formic acid) in 20 min at a flow rate of 5 mL/min. The radiolabeled product was collected (RT; 13.4 min for AJ662, 13.4 min for AJ698, 12.5 min for AJ699) with decay-corrected radiochemical yield of > 95%. The desired radiolabeled fraction was concentrated under a stream of N_2_ at 60°C, formulated in 0.1% Tween^20^ in saline, and used for *In vitro* and *In vivo* studies. The whole radiolabeling process was completed in approximately 90 min. Quality control was performed on the same HPLC system using the same HPLC gradient as described above.

### Method for Lu-177 radiolabeling

The ^177^Lu was obtained from National Isotope Development Center (NIDC) and dissolved in 0.1 M NaOAc (pH 5.0). To a microcentrifuge vial (1.5 mL) containing 200 μL of 0.1 M NaOAc buffer (pH = 5), 10 μL of 1 M of ascorbic acid and 40 µg of AJ662 (around 20 nmoles), 8.1 mCi of [^177^Lu]Lu(OAc)_3_ was added. The reaction mixture was incubated for 60 min at 90 °C in a temperature-controlled heating block and purified on a RP-HPLC system using a semi-preparative C-18 Luna column (5 mm, 10 x 250 mm Phenomenex, Torrance, CA). The HPLC conditions were gradient elution starting with 5% acetonitrile: water (0.1% trifluoroacetic acid) and reaching 95% acetonitrile: water (0.1% trifluoroacetic acid) in 20 min at a flow rate of 5 mL/min. The radiolabeled product was collected at the retention time of 13.4 min with decay-corrected radiochemical yield of around 68 %. The desired radiolabeled fraction was concentrated under a stream of N_2_ at 60 °C, formulated in 0.1% Tween^20^ in saline, and used for *In vitro* and *In vivo* studies. The whole radiolabeling process was completed in approximately 90 min. Quality control was performed on the same HPLC system using the same HPLC gradient as described above.

### Cell Culture

SCaBER, and BFTC909 cell lines were purchased from ATCC and cultured in DMEM media and were supplemented with 10% FBS and 1% P/S antibiotic. These cell lines were routinely tested for Mycoplasma. Freshly thawed cells were cultured for less than 3 months or authenticated at JHU Genomics Core.

### Flow Cytometry Analysis for PD-L1 Expression

SCaBER and BFTC909 cells are adherent and were detached using enzyme-free cell dissociation buffer (Thermo Fisher Scientific, Waltham, MA). To evaluate PD-L1 surface expression, 1×10^6^ cells were incubated in 100 μL FACS buffer (PBS with 0.1% FBS and 2 mmol/L ethylenediaminetetraacetic acid) with anti-PD-L1 antibody (clone: MIH1 BD # 558065) for 30 minutes at 4°C on ice. After washing and resuspending in FACS buffer, cell sorting was done on a BD Accuri C6 plus flow cytometer and data was analyzed using FlowJo software.

### General Procedure For *In Vitro* Cellular Binding Assays

*In vitro* binding assays were conducted to determine the binding of radiolabeled peptides to cells. Approximately 1 μCi of radiotracer (F-18 or Ga-68 or Lu-177 labelled peptides) were incubated with around 1×10^6^ cells for 60 min at 4 °C. After incubation, cells were washed three times with ice-cold PBS containing 0.1% tween-20 and bound radioactivity on cell pellets were counted on an automated gamma counter (1282 Compugamma CS, Pharmacia/LKB Nuclear, Inc., Gaithersburg, MD). Blocking was performed with 2 µM of non-radiolabeled precursor to demonstrate receptor-specific cellular binding of a tracer. All cellular uptake studies were performed in quadruplicate for each cell line and repeated three times.

### Tumor Models

Animal studies were performed under Johns Hopkins University ACUC-approved protocols. Male NSG mice (5-6 weeks old) were used to establish xenografts by administering 3-4 million cells (SCaBER in left and BFTC909 in right,) subcutaneously in upper flanks to form tumor models within 3-4 weeks. Imaging or biodistribution studies (n=3-5) were conducted on mice with tumor volumes of 100-200 mm^3^.

### General Method For PET-CT Imaging of Mouse Xenografts with Radiotracers

Mice bearing flank tumors were injected intravenously with ^68^Ga or ^18^F labelled radiotracers and PET images were acquired 60 and 120 minutes after injection of the radiotracer. Images were acquired in 2 bed positions for a total of 10 minutes using an ARGUS small-animal PET/CT scanner. Images were reconstructed using 2D-OSEM and corrected for radioactive decay and dead time. Image fusion, visualization, and 3D rendering were accomplished using Amira 2020.3.1.

### *Ex Vivo* Biodistribution

Mice with 100-200 mm^3^ tumor volume were used for *ex vivo* biodistribution studies. For radiotracer pharmacokinetics studies, mice with SCaBER, BFTC909 and MOLP8 tumor xenograft received ∼80 µCi (2.96 MBq) of radiotracers and were sacrificed at pre-determined time points (2, 30, 60 and 120 min). The selected tissues were harvested, weighed, counted, and their % IA g^-1^ values calculated for biodistribution analysis. The tissues included blood, muscle, femur, both tumors, Lungs, heart, pancreas, spleen, liver, small intestine, large intestine, stomach, kidneys, bladder and brain.

### PD-L1 Immunohistochemistry of Tumor Sections

After the imaging study, mice were humanely euthanized, and their tumors were harvested and fixed in 4% paraformaldehyde and sent to oncology tissue services (OTS) at JHU. Tissue sections of 4 µm thickness were prepared from paraffin-embedded tumors. These tumor sections (BFTC909 and SCaBER) were evaluated for PD-L1 expression by IHC as reported previously^26^. Briefly, 4–5-μm tumor sections were treated with 3% H_2_O_2_ for 10 minutes, blocked with 5% goat serum for 1 hour, and then incubated with a primary anti-human PD-L1 antibody (#13684, Cell Signaling Technology) at 1:500 dilution at 4°C overnight. Subsequently, using Dako CSAII Biotin-free Tyramide Signal Amplification System Kit, slides were incubated with secondary antibody and amplification reagent. Final staining was carried out by adding 3,3’-Diaminobenzidine. After washing, the slides were counterstained with Mayer’s Hematoxylin for 1 min, dehydrated using alcohol and xylene, and then cover slipped. ^1^

### In Vitro and In Vivo Stability Study

For in vitro stability study, radiotracers (∼100 µCi) were aliquoted at specified time points from stock solutions maintained at room temperature in saline containing 0.1% Tween-20. These aliquots were analyzed using RP-HPLC equipped with a radioactivity detector. For In vivo stability, radiotracers (around 300 µCi) were injected intravenously into the naïve mice. Urine and blood were collected at 60 and 120 min after radiotracer injection. The blood samples were centrifuged at 13,000 rpm for 15 min to separate blood capsules and serum was collected. This serum and urine samples were diluted with ice-cold acetonitrile (1:1 v/v), centrifuged at 13,000 rpm for 5 min at 4 °C and the supernatant were collected and filtered with 0.22 μm syringe filter.40,41 These samples were analyzed using RP-HPLC connected to a radio-detector, allowing us to distinguish between degraded and intact radiolabeled peptide. The percentage of intact radiolabeled peptide in the urine and serum was quantified by calculating the ratio of the AUC for the intact peak in the radioactivity detector to that of its degradation products (free radioactivity).

### Statistical Analysis

All statistical analyses were performed using Prism 10.6 Software (GraphPad Software, La Jolla, CA). Unpaired Student’s *t test* and one- or two-way ANOVA were utilized for column, multiple column, and grouped analyses, respectively. Statistical significance was set at ns, *P* ≥ 0.05; *, *P* ≤ 0.05; **, *P* ≤ 0.01; ***, *P* ≤ 0.001; ****, *P* ≤ 0.0001.

## Acknowledgements

The authors thank Ms. Xiaoju Yang for conducting PET-CT studies in MRB molecular imaging service center and cancer functional imaging core. The authors also thank Drs. Aykut Üren and Purushottam Tiwari at Georgetown University for performing SPR experiments.

## Author Contributions

A.K.S.: Conceptualization, methodology, data curation, formal analysis, writing-original draft, writing-review and editing. A.M.: Methodology, data curation, writing-review and editing. K.G.: Methodology, data curation, writing-review and editing. S.N.: Funding acquisition, conceptualization, methodology, data curation, formal analysis, writing-original draft, writing-review and editing.

## Data Availability

All study data are included in the article and/or supplementary information. Additional data are available upon request from the corresponding author.

## References

1. Bodei, L., Herrmann, K., Schoder, H., Scott, A.M. & Lewis, J.S. Radiotheranostics in oncology: current challenges and emerging opportunities. Nat Rev Clin Oncol 19, 534–550 (2022).

2. Primac, I., Tabury, K., Tasdogan, A., Baatout, S. & Herrmann, K. The molecular blueprint of targeted radionuclide therapy. Nat Rev Clin Oncol 22, 869–894 (2025).

3. Halder, R. & Ritter, T. (18)F-Fluorination: Challenge and Opportunity for Organic Chemists. J Org Chem 86, 13873–13884 (2021).

4. Ma, M.T., Boros, E. & Wilson, J.J. The Inorganic Chemistry of Radiopharmaceuticals. Inorganic chemistry 62, 20537–20538 (2023).

5. McBride, W.J., D’Souza, C.A., Sharkey, R.M. & Goldenberg, D.M. The radiolabeling of proteins by the [18F]AlF method. Appl Radiat Isot 70, 200–204 (2012).

6. Perrin, D.M. [(18)F]-Organotrifluoroborates as Radioprosthetic Groups for PET Imaging: From Design Principles to Preclinical Applications. Acc Chem Res 49, 1333–1343 (2016).

7. Huang, S., et al. Comparison of 18F-based PSMA radiotracers with [68Ga]Ga-PSMA-11 in PET/CT imaging of prostate cancer—a systematic review and meta-analysis. Prostate Cancer and Prostatic Diseases 27, 654–664 (2024).

8. Cerci, J.J., et al. Diagnostic Performance and Clinical Impact of ^68^Ga-PSMA-11 PET/CT Imaging in Early Relapsed Prostate Cancer After Radical Therapy: A Prospective Multicenter Study (IAEA-PSMA Study). Journal of Nuclear Medicine 63, 240–247 (2022).

9. Rowe, S.P., Gorin, M.A. & Pomper, M.G. Imaging of Prostate-Specific Membrane Antigen Using [18F]DCFPyL. PET Clinics 12, 289–296 (2017).

10. Ulaner, G.A., et al. 18F-DCFPyL PET/CT for Initially Diagnosed and Biochemically Recurrent Prostate Cancer: Prospective Trial with Pathologic Confirmation. Radiology 305, 419–428 (2022).

11. Jones, L.H. & Kelly, J.W. Structure-based design and analysis of SuFEx chemical probes. RSC Med Chem 11, 10–17 (2020).

12. Rojas, J.J. & Bull, J.A. Unconventional reactivity of sulfonyl fluorides. Trends in Chemistry 7, 124–133 (2025).

13. Zheng, Q., et al. Sulfur [(18)F]Fluoride Exchange Click Chemistry Enabled Ultrafast Late-Stage Radiosynthesis. J Am Chem Soc 143, 3753–3763 (2021).

14. Wang, Z., et al. Design, synthesis and evaluation of novel prostate-specific membrane antigen-targeted aryl [(18)F]fluorosulfate PET tracers. Bioorg Med Chem 106, 117753 (2024).

15. Craig, A., et al. Preparation of (18)F-Labeled Tracers Targeting Fibroblast Activation Protein via Sulfur [(18)F]Fluoride Exchange Reaction. Pharmaceutics 15(2023).

16. Huang, Y., et al. Design of Novel (18)F-Labeled Amino Acid Tracers Using Sulfur (18)F-Fluoride Exchange Click Chemistry. ACS medicinal chemistry letters 15, 294–301 (2024).

17. Deiser, S., et al. (SiFA)SeFe: A Hydrophilic Silicon-Based Fluoride Acceptor Enabling Versatile Peptidic Radiohybrid Tracers. J Med Chem 67, 14077–14094 (2024).

18. Kumar, D., et al. Pharmacodynamic measures within tumors expose differential activity of PD(L)-1 antibody therapeutics. Proceedings of the National Academy of Sciences 118, e2107982118 (2021).

19. Kumar, D., et al. Peptide-based PET quantifies target engagement of PD-L1 therapeutics. The Journal of Clinical Investigation 129, 616–630 (2018).

20. Mishra, A., et al. Gallium-68–labeled Peptide PET Quantifies Tumor Exposure of PD-L1 Therapeutics. Clinical Cancer Research 29, 581–591 (2023).

21. Bocuzzi, V., et al. CD38 as theranostic target in oncology. Journal of Translational Medicine 22, 998 (2024).

22. Sharma, A.K., et al. CD38-Specific Gallium-68 Labeled Peptide Radiotracer Enables Pharmacodynamic Monitoring in Multiple Myeloma with PET. Advanced Science 11, 2308617 (2024).

23. Veryser, C., Demaerel, J., Bieliu̅nas, V., Gilles, P. & De Borggraeve, W.M. Ex Situ Generation of Sulfuryl Fluoride for the Synthesis of Aryl Fluorosulfates. Organic Letters 19, 5244–5247 (2017).

24. Chen, W., et al. Synthesis of Sulfotyrosine-Containing Peptides by Incorporating Fluorosulfated Tyrosine Using an Fmoc-Based Solid-Phase Strategy. Angewandte Chemie International Edition 55, 1835–1838 (2016).

25. Sharma, A.K., et al. EphA2-targeted alpha-particle theranostics for enhancing PDAC treatment. Theranostics 15, 4229–4246 (2025).

26. Chatterjee, S., et al. A humanized antibody for imaging immune checkpoint ligand PD-L1 expression in tumors. Oncotarget 7, 10215–10227 (2016).

