## Supplementary material for "SuFEx and macrocyclic chelation define orthogonal reactivity domains enabling isotopically versatile radiopharmaceuticals from a single peptide precursor": Supplemetal File

for

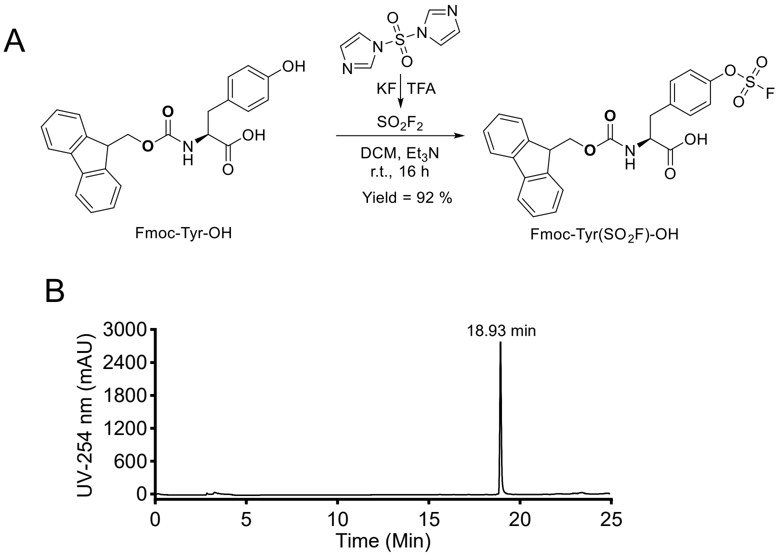


**Figure S1. Synthesis and characterization of Fmoc-Tyr($SO_2F$)-OH. A)** Synthetic scheme of Fmoc-Tyr($SO_2F$)-OH. Fmoc-Tyr-OH was reacted with ex situ generated $SO_2F_2$ in the presence of triethylamine ($Et_3N$) using dichloromethane (DCM) as the solvent at room temperature overnight, affording Fmoc-Tyr($SO_2F$)-OH in >90% yield. **B)** HPLC trace of Fmoc-Tyr($SO_2F$)-OH demonstrating its high purity.

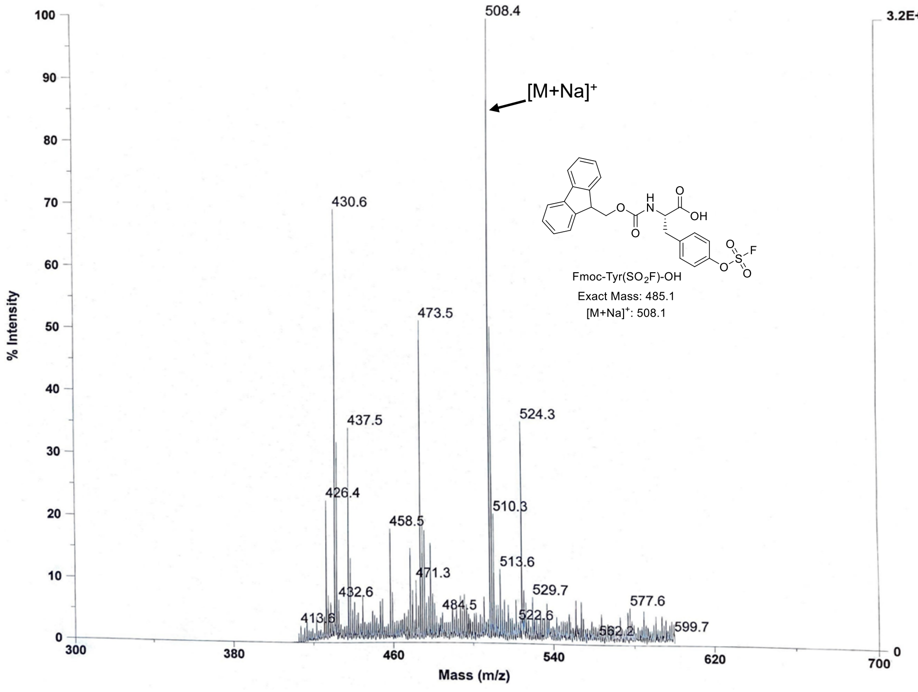


**Figure S2.** Characterization of Fmoc-Tyr($SO_2F$)-OH by MALDI-TOF mass spectrometry

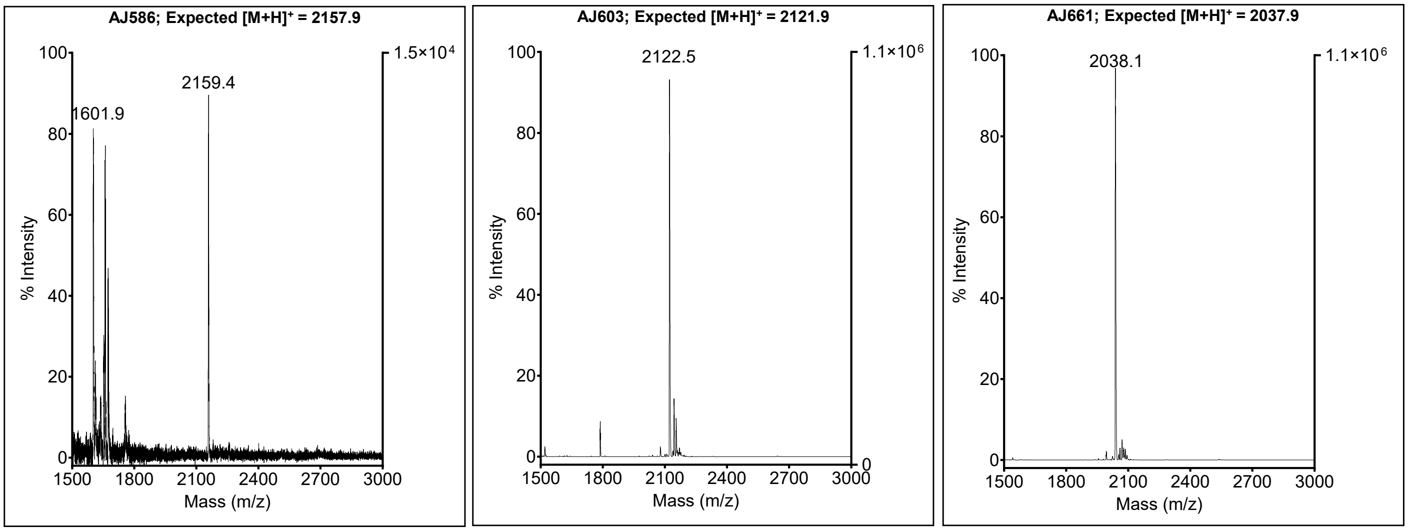


**Figure S3.** Characterization of AJ586, AJ603, and AJ661 using MALDI-TOF Mass spectrometry

**Table S1.** Ex vivo biodistribution of $[^{18}F]$AJ603 in BFTC909 and SCaBER tumor-bearing NSG mice collected at 2, 60, 120, and 240 min post-injection

| | Percentage incubated activity per gram (%IA/g) in Mean±SEM (n=4) | | | |
|---|---|---|---|---|
| Tissues | 2 min | 60 min | 120 min | 240 min |
| Blood | 36.36±2.65 | 13.91±0.57 | 8.79±0.47 | 5.94±0.43 |
| Muscle | 0.64±0.02 | 0.79±0.06 | 0.67±0.06 | 0.41±0.01 |
| Femur | 2.01±0.19 | 1.41±0.04 | 1.38±0.04 | 1.64±0.11 |
| SCaBER | 1.28±0.13 | 2.42±0.10 | 1.94±0.13 | 1.96±0.18 |
| BFTC909 | 0.83±0.05 | 3.93±0.34 | 4.01±0.51 | 4.98±0.14 |
| Lung | 18.24±1.75 | 6.92±0.32 | 4.40±0.20 | 3.46±0.28 |
| Heart | 9.55±0.25 | 3.86±0.26 | 2.72±0.10 | 1.63±0.11 |
| Pancreas | 3.06±0.16 | 1.29±0.06 | 0.89±0.07 | 1.44±0.37 |
| Spleen | 0.99±0.37 | 2.21±0.34 | 1.68±0.06 | 1.43±0.09 |
| Liver | 15.72±1.46 | 15.30±0.59 | 9.81±1.32 | 8.33±0.59 |
| Small intestine | 1.47±0.19 | 3.79±1.48 | 11.72±2.36 | 10.06±4.08 |
| Large intestine | 0.37±0.05 | 0.40±0.05 | 0.61±0.15 | 9.74±1.01 |
| Stomach | 0.59±0.02 | 0.86±0.06 | 0.81±0.31 | 1.47±0.24 |
| Kidneys | 8.94±0.42 | 13.02±1.74 | 12.95±1.91 | 10.31±0.95 |
| Bladder | 1.24±0.18 | 3.89±0.55 | 12.98±2.08 | 18.09±4.36 |
| Brain | 0.83±0.12 | 0.32±0.06 | 0.15±0.02 | 0.18±0.01 |

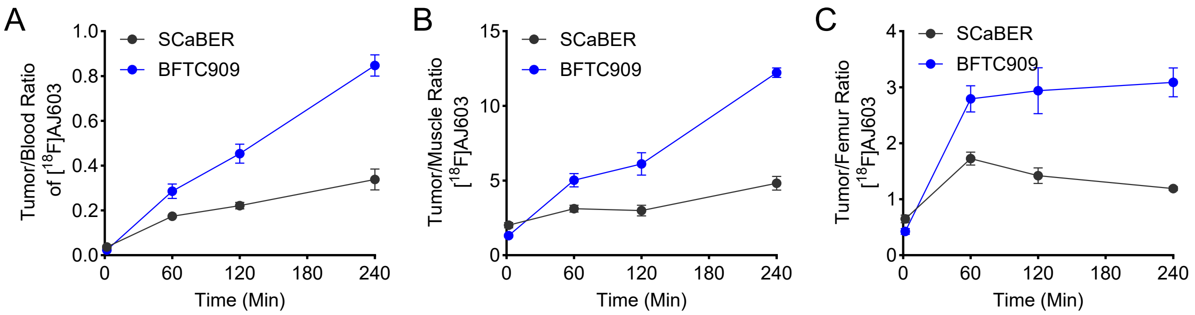


**Figure S4. In vivo time-activity curves of tumor-to-tissue ratios in BFTC909 and SCaBER xenograft models.** Time-activity curves showing **A)** tumor-to-blood, **B)** tumor-to-muscle and **C)** tumor-to-femur ratios following administration of [$^{18}$F]AJ603 in BFTC909 and SCaBER tumor-bearing mice. Data were derived from ex vivo biodistribution studies up to 240 min post-injection and are presented as mean ± SEM (n = 4 per time point).

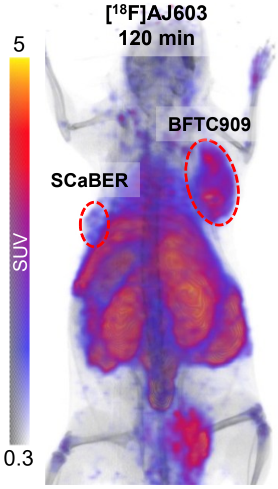


**Figure S5. PET imaging with [$^{18}$F]AJ603 in SCaBER and BFTC909 tumor-bearing mice.** Whole-body 3D rendered PET images illustrating overall radiotracer distribution. High [$^{18}$F]AJ603 uptake was observed in BFTC909 tumors, while SCaBER tumors showed minimal accumulation, demonstrating PD-L1–specific binding of the tracer.

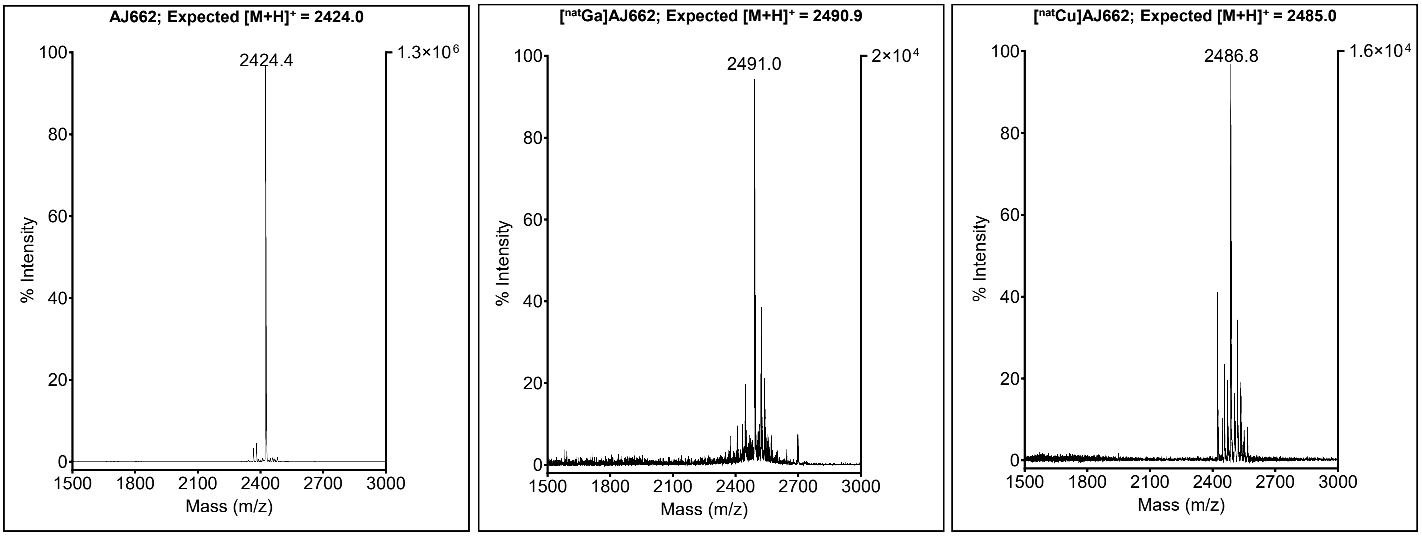


**Figure S6.** Characterization of AJ662, [$^{nat}$Ga]AJ662 and [$^{nat}$Cu]AJ662 using MALDI-TOF Mass spectrometry

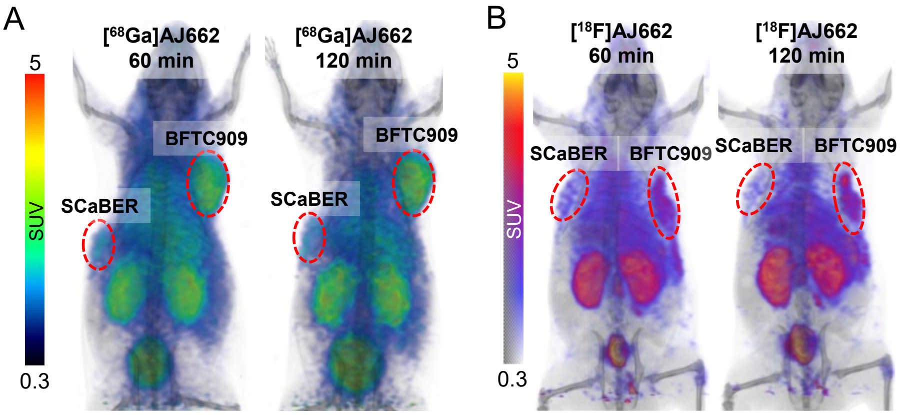


**Figure S7. Whole-body 3D rendered PET images of PD-L1–targeted radiotracers in SCaBER and BFTC909 tumor-bearing mice.** Whole-body 3D rendered PET images illustrating the biodistribution of radiotracers in vivo. **A)** PET images acquired at 60 and 120 min post-injection with [$^{68}$Ga]AJ662 **B)** PET images acquired at 60 and 120 min post-injection with [$^{18}$F]AJ662. High radiotracer accumulation was observed in BFTC909 tumors, whereas SCaBER tumors showed minimal uptake, confirming PD-L1–specific binding of these tracers.

**Table S2.** Ex vivo biodistribution of [$^{68}$Ga]AJ662 in BFTC909 and SCaBER tumor-bearing NSG mice collected at 2, 30, 60, and 120 min post-injection

| | Percentage incubated activity per gram (%IA/g) in Mean±SEM (n=4) | | | | |
|---|---|---|---|---|---|
| Tissues | 2 min | 30 min | 60 min | 120 min | 120-blocking |
| Blood | 52.82±7.21 | 13.14±2.08 | 6.58±0.37 | 2.08±0.07 | 2.81±0.54 |
| Muscle | 1.19±0.20 | 1.41±0.08 | 0.89±0.08 | 0.55±0.04 | 0.64±0.01 |
| Femur | 2.61±0.47 | 1.68±0.12 | 1.21±0.04 | 0.89±0.11 | 0.88±0.09 |
| SCaBER | 1.55±0.15 | 3.70±0.16 | 2.49±0.50 | 1.93±0.25 | 2.01±0.04 |
| BFTC909 | 1.84±0.27 | 6.32±0.39 | 8.93±0.14 | 7.65±1.08 | 2.68±0.42 |
| Lungs | 22.61±0.99 | 6.60±0.47 | 4.05±0.18 | 1.78±0.19 | 2.18±0.33 |
| Hearts | 13.73±0.73 | 4.46±0.29 | 2.59±0.15 | 1.30±0.09 | 1.36±0.15 |
| Pancreas | 3.89±0.88 | 1.53±0.16 | 1.12±0.06 | 0.76±0.09 | 1.65±0.75 |
| Spleen | 2.13±1.13 | 2.10±0.17 | 2.00±0.16 | 1.68±0.30 | 2.63±0.72 |
| Liver | 11.74±1.83 | 4.74±0.18 | 4.11±0.16 | 2.60±0.11 | 2.54±0.79 |
| Small intestine | 3.64±1.53 | 1.84±0.08 | 1.79±0.08 | 1.74±0.33 | 2.31±0.78 |
| Large intestine | 1.31±0.54 | 0.39±0.02 | 0.39±0.03 | 0.45±0.04 | 0.54±0.09 |
| Stomach | 2.03±0.65 | 0.62±0.11 | 0.67±0.02 | 0.42±0.03 | 0.75±0.35 |
| Kidneys | 17.60±1.50 | 13.00±1.04 | 12.05±0.40 | 13.16±1.44 | 13.28±1.21 |
| Bladder | 2.07±0.45 | 27.85±14.77 | 31.57±7.22 | 47.34±18.94 | 51.07±27.01 |
| Brain | 1.23±0.38 | 0.39±0.07 | 0.21±0.03 | 0.08±0.02 | 0.12±0.01 |

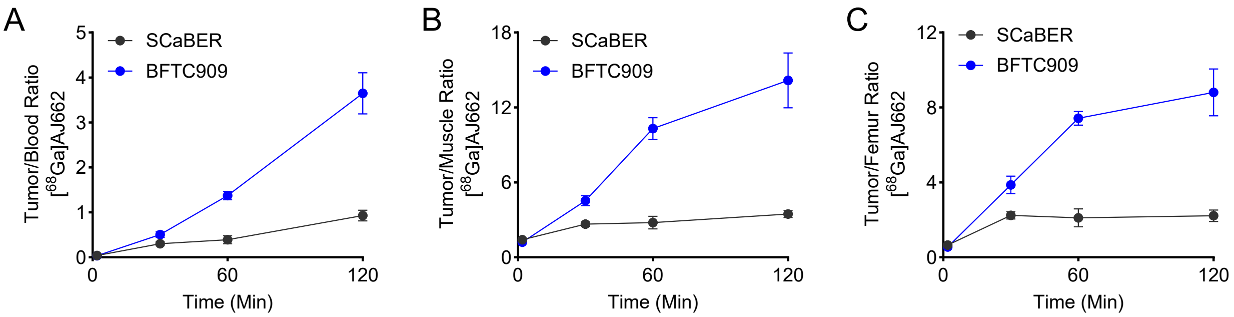


**Figure S8 . In vivo time-activity curves of tumor-to-tissue ratios in BFTC909 and SCaBER xenograft models.** Time-activity curves showing **A)** tumor-to-blood, **B)** tumor-to-muscle and **C)** tumor-to-femur ratios following administration of [$^{68}$Ga]AJ662 in BFTC909 and SCaBER tumor-bearing mice. Data were obtained from ex vivo biodistribution studies up to 120 min post-injection and are presented as mean ± SEM (n = 4 per time point).

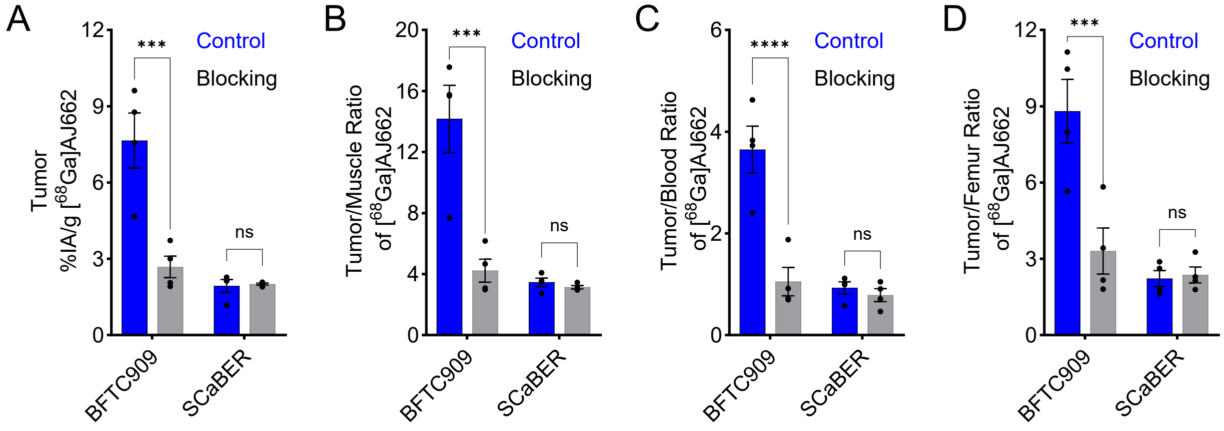


**Figure S9. In vivo PD-L1 specificity assessment of [$^{68}$Ga]AJ662 in BFTC909 and SCaBER xenograft models. A)** Tumor radioactivity uptake (%IA/g), **B)** tumor-to-muscle, **C)** tumor-to-blood **D)** tumor-to-femur ratios were determined from ex vivo biodistribution data at 120 min post-injection of [$^{68}$Ga]AJ662. For blocking studies, mice were treated with an excess dose of AJ662 (1 mg/kg; ~50 µg per animal) administered 30 min prior to radiotracer injection. Data are presented as mean ± SEM (n = 4 per group). Statistical significance was determined using two-way ANOVA; ns, P ≥ 0.05; ***, P ≤ 0.001; ****, P ≤ 0.0001.

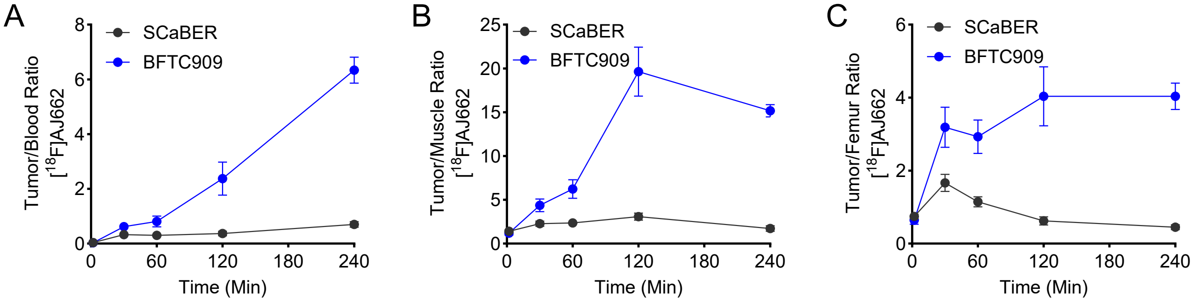


**Figure S10. In vivo time-activity curves of tumor-to-tissue ratios in BFTC909 and SCaBER xenograft models.** Time-activity curves showing **A)** tumor-to-blood, **B)** tumor-to-muscle and **C)** tumor-to-femur ratios following administration of [$^{18}$F]AJ662 in BFTC909 and SCaBER tumor-bearing mice. Data were derived from ex vivo biodistribution studies up to 120 min post-injection and are presented as mean ± SEM (n = 4 per time point).

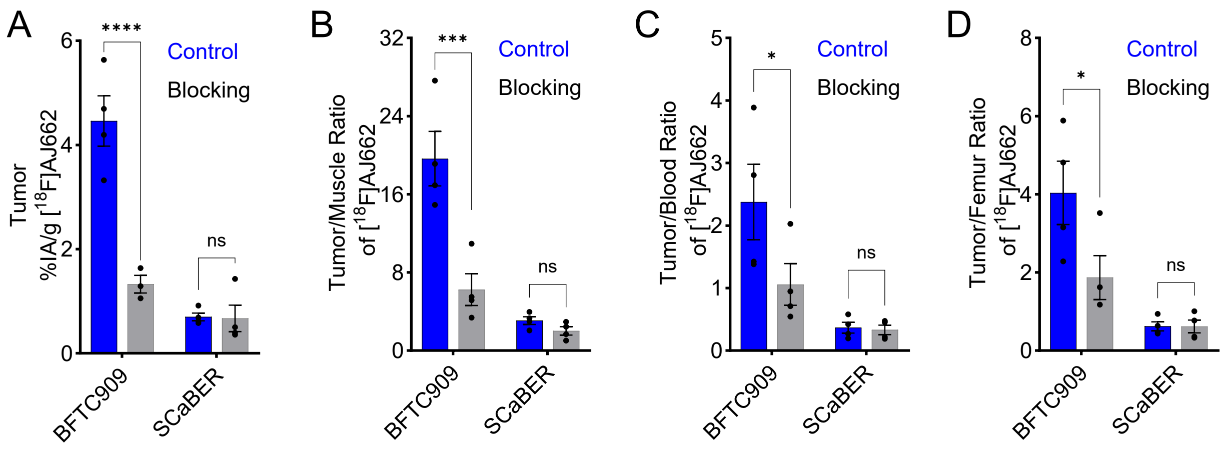


**Figure S11. In vivo PD-L1 specificity assessment of [$^{18}$F]AJ662 in BFTC909 and SCaBER xenograft models. A)** Tumor radioactivity uptake (%IA/g), **B)** tumor-to-muscle, **C)** tumor-to-blood **D)** tumor-to-femur ratios were determined from ex vivo biodistribution data at 120 min post-injection of [$^{18}$F]AJ662. For blocking studies, mice were treated with an excess dose of AJ662 (1 mg/kg; ~50 µg per animal) administered 30 min prior to radiotracer injection. Data are presented as mean ± SEM (n = 4 per group). Statistical significance was determined using two-way ANOVA; ns, P ≥ 0.05; ***, P ≤ 0.001; ****, P ≤ 0.0001.

**Table S3.** Ex vivo biodistribution of [$^{18}$F]AJ662 in BFTC909 and SCaBER tumor-bearing NSG mice collected at 2, 30, 60, and 120 min post-injection

| | Percentage incubated activity per gram (%IA/g) in Mean±SEM (n=4) | | | | |
|---|---|---|---|---|---|
| Tissues | 2 min | 30 min | 60 min | 120 min | 120 with blocking |
| Blood | 37.34±3.96 | 8.36±0.41 | 7.83±2.07 | 2.25±0.55 | 1.96±0.39 |
| Muscle | 0.98±0.13 | 1.30±0.27 | 0.92±0.18 | 0.23±0.03 | 0.32±0.06 |
| Femur | 1.79±0.13 | 1.77±0.35 | 1.89±0.28 | 1.18±0.13 | 1.06±0.16 |
| SCaBER | 1.31±0.02 | 2.73±0.14 | 2.15±0.42 | 0.70±0.08 | 0.67±0.25 |
| BFTC909 | 1.11±0.12 | 5.13±0.22 | 5.17±0.39 | 4.46±0.48 | 1.95±0.64 |
| Lungs | 16.44±1.73 | 5.09±0.20 | 4.54±1.21 | 1.08±0.07 | 1.36±0.19 |
| Hearts | 10.89±1.60 | 2.50±0.13 | 2.20±0.53 | 0.51±0.03 | 0.76±0.14 |
| Pancreas | 4.19±0.66 | 1.50±0.17 | 1.92±0.62 | 1.03±0.60 | 0.87±0.39 |
| Spleen | 1.27±0.20 | 2.02±0.17 | 1.69±0.50 | 0.53±0.03 | 0.92±0.26 |
| Liver | 9.10±1.12 | 3.61±0.25 | 3.72±0.96 | 1.45±0.15 | 1.58±0.21 |
| Small intestine | 3.06±1.00 | 1.36±0.33 | 5.58±2.03 | 1.59±0.45 | 6.64±3.30 |
| Large intestine | 1.69±0.82 | 0.45±0.08 | 0.49±0.15 | 0.89±0.22 | 0.52±0.14 |
| Stomach | 2.63±1.11 | 0.19±0.04 | 1.01±0.23 | 1.12±0.66 | 0.52±0.35 |

| | | | | | |
|---|---|---|---|---|---|
| Kidneys | 12.11±1.22 | 11.84±0.56 | 14.92±2.87 | 4.46±1.14 | 5.42±1.49 |
| Bladder | 2.45±0.40 | 34.56±16.60 | 11.98±3.73 | 24.77±15.38 | 60.87±27.57 |
| Brain | 0.84±0.06 | 0.18±0.02 | 0.20±0.04 | 0.05±0.01 | 0.07±0.03 |

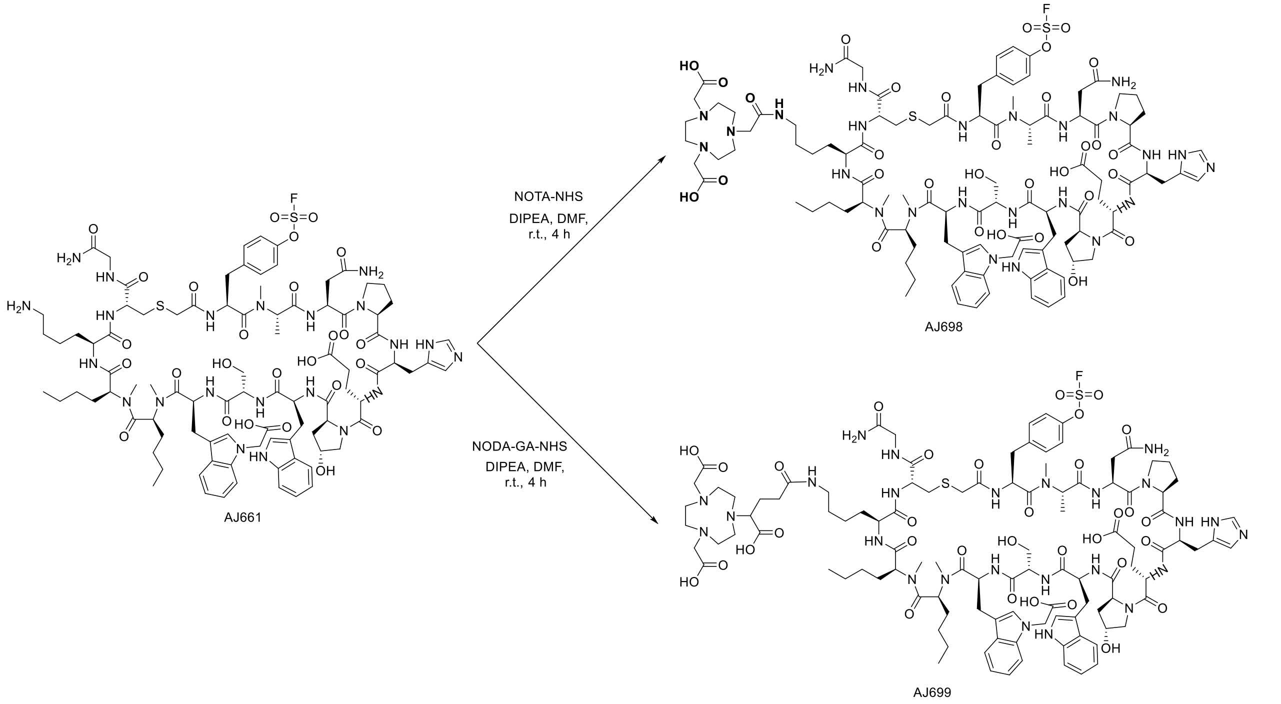


**Scheme S1**. Synthesis of AJ698 and AJ699, PD-L1 binding peptide containing NOTA and NODAGA chelator. These peptides were synthesized by conjugating AJ661 with NOTA-NHS and NODAGA NHS ester, respectively, in the presence of DIPEA as a base and DMF as solvent at room temperature.

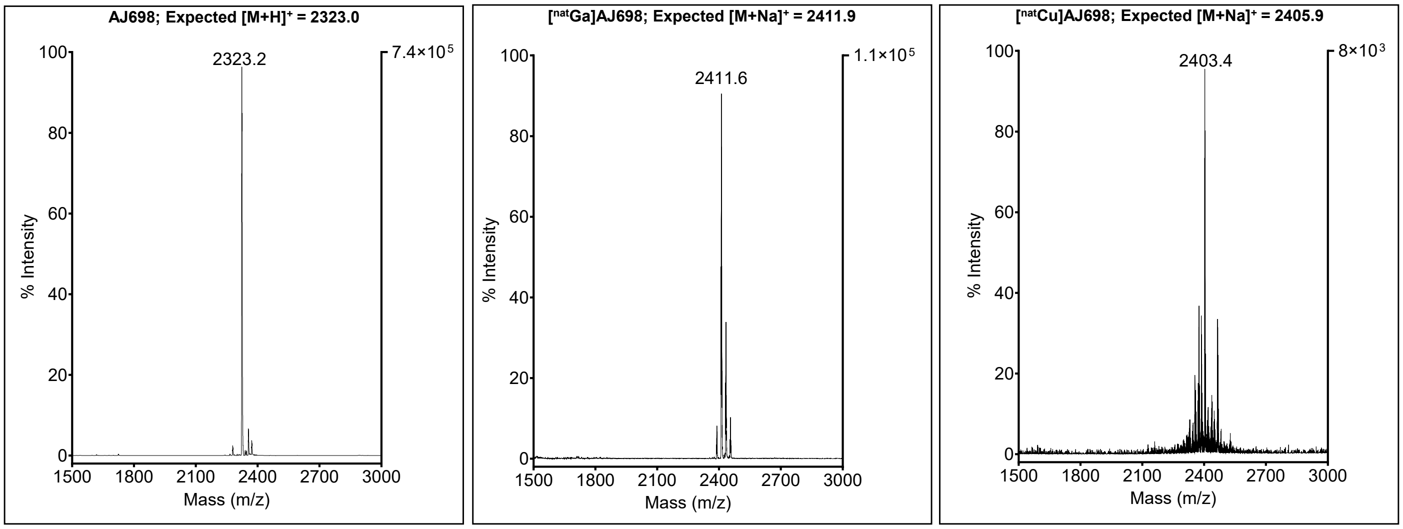


**Figure S12.** Characterization of AJ698, [$^{nat}$Ga]AJ698 and [$^{nat}$Cu]AJ698 using MALDI-TOF Mass spectrometry

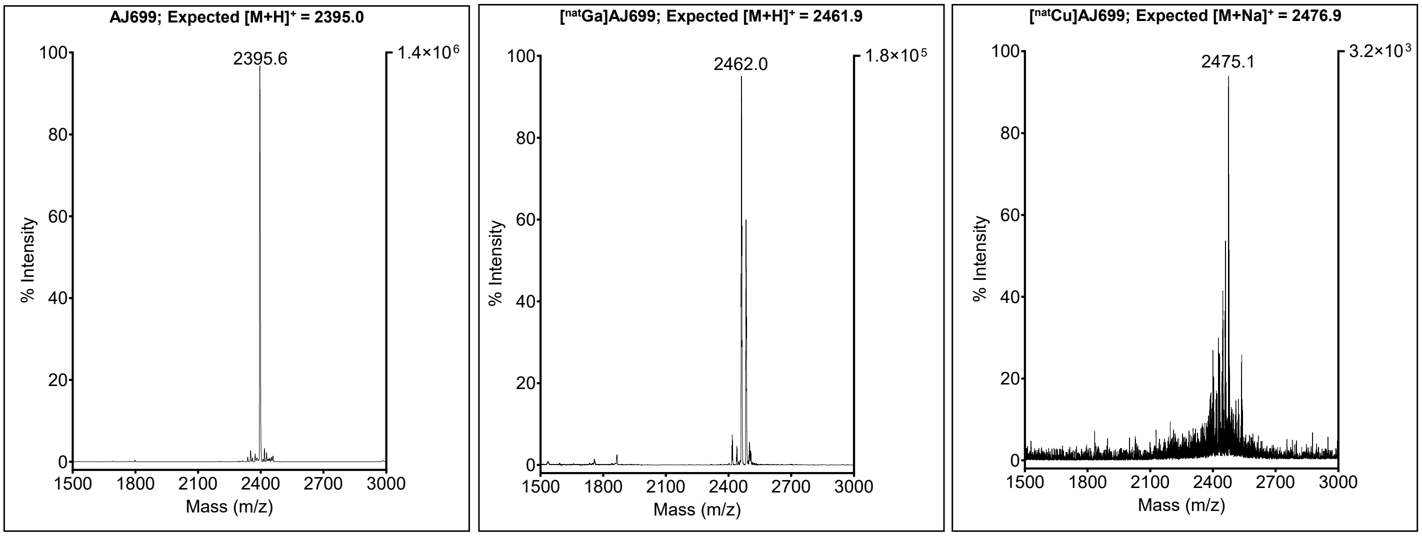


**Figure S13.** Characterization of AJ699, [$^{nat}$Ga]AJ699 and [$^{nat}$Cu]AJ699 using MALDI-TOF Mass spectrometry

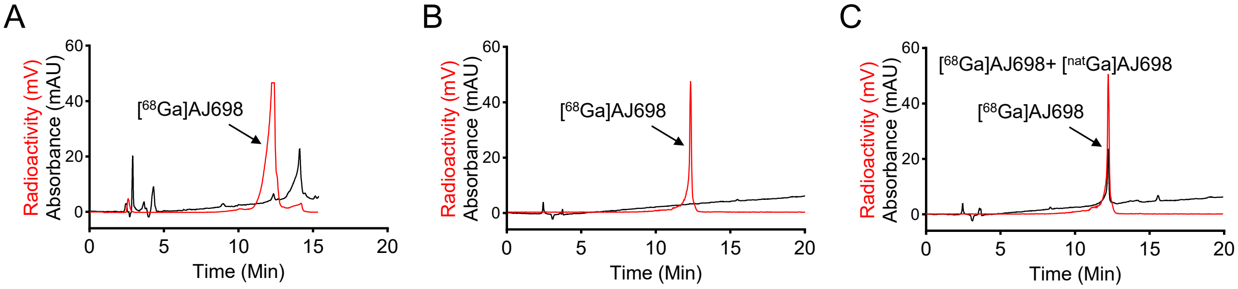


**Figure S14. Radiolabeling and quality control of [$^{68}$Ga]AJ698. A)** HPLC chromatogram of radiotracer purification showing a decay-corrected radiochemical yield of >95%, confirming efficient labeling of AJ698. **B)** HPLC chromatogram representing quality control analysis of [$^{68}$Ga]AJ698 prior to further biological evaluation. **C)** HPLC chromatogram confirming the chemical identity of [$^{68}$Ga]AJ698 by co-elution with its non-radioactive counterpart [$^{nat}$Ga]AJ698. The identical retention times and peak profiles confirm successful and specific incorporation of $^{68}$Ga into the AJ698 peptide.

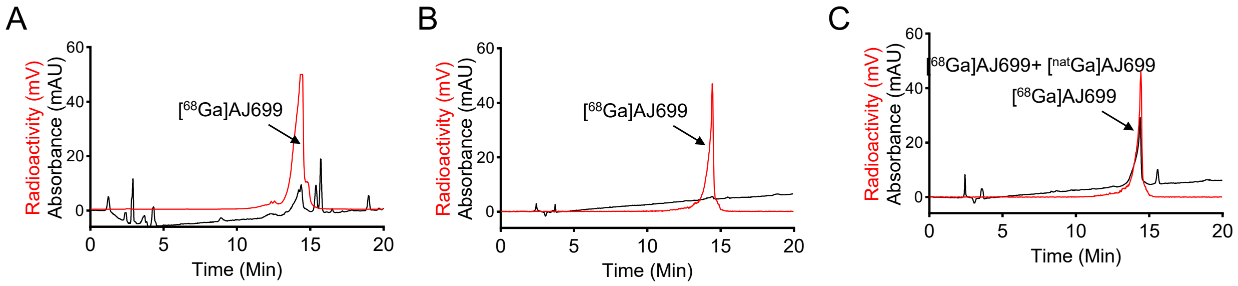


**Figure S15. Radiolabeling and quality control of [$^{68}$Ga]AJ699. A)** HPLC chromatogram of radiotracer purification showing a decay-corrected radiochemical yield of >95%, confirming efficient labeling of AJ699. **B)** HPLC chromatogram representing quality control analysis of [$^{68}$Ga]AJ699 prior to further biological evaluation. **C)** HPLC chromatogram confirming the chemical identity of [$^{68}$Ga]AJ699 by co-elution with its non-radioactive counterpart [$^{nat}$Ga]AJ699. The identical retention times and peak profiles confirm successful and specific incorporation of $^{68}$Ga into the AJ699 peptide.

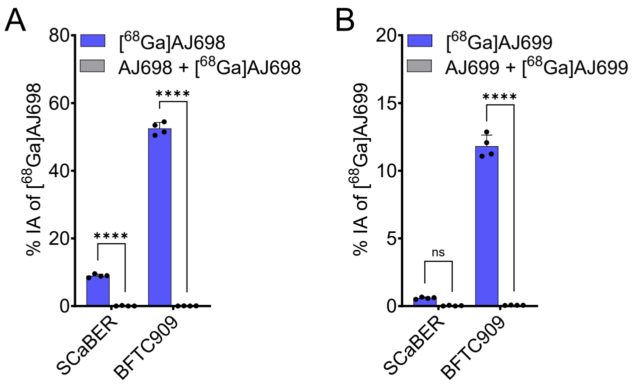


**Figure S16. In vitro evaluation of [$^{68}$Ga]AJ698 and [$^{68}$Ga]AJ699 in bladder cancer cells. A)** Cellular binding of [$^{68}$Ga]AJ698 (%IA) in bladder cancer cells following incubation with 1 µCi of radiotracer at 4 °C for 1 h. (B) Cellular binding of [$^{68}$Ga]AJ699 under identical conditions. Co-incubation with 1 µM of the corresponding non-radioactive compounds (blocking dose) significantly reduced radiotracer uptake, confirming PD-L1–specific binding.

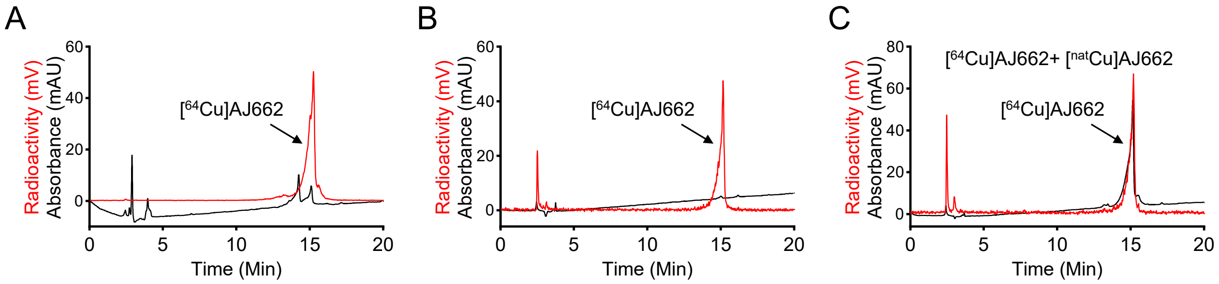


**Figure S17. Radiolabeling and quality control of [$^{64}$Cu]AJ662. A)** HPLC chromatogram of radiotracer purification showing a decay-corrected radiochemical yield of >95%, confirming efficient labeling of AJ662. **B)** HPLC chromatogram representing quality control analysis of [$^{64}$Cu]AJ662 prior to further biological evaluation. **C)** HPLC chromatogram confirming the chemical identity of [$^{64}$Cu]AJ662 by co-elution with its non-radioactive counterpart [$^{nat}$Cu]AJ662. The identical retention times and peak profiles confirm successful and specific incorporation of $^{64}$Cu into the AJ662 peptide.

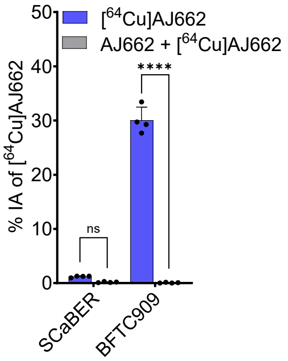


**Figure S18. In vitro binding** (%IA) **of [$^{64}$Cu]AJ662 in bladder cancer cells**. Cells were incubated with 0.2 µCi of radiotracer at 4 °C for 1 h. Co-incubation with 1 µM of the corresponding non-radioactive peptide (blocking dose) significantly reduced radiotracer uptake, confirming PD-L1–specific binding.

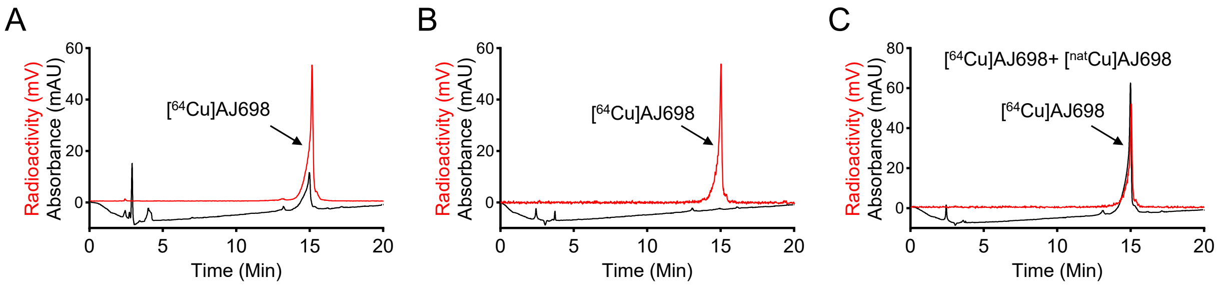


**Figure S19. Radiolabeling and quality control of [$^{64}$Cu]AJ698. A)** HPLC chromatogram of radiotracer purification showing a decay-corrected radiochemical yield of >95%, confirming efficient labeling of AJ698. **B)** HPLC chromatogram representing quality control analysis of [$^{64}$Cu]AJ698 prior to further biological evaluation. **C)** HPLC chromatogram confirming the chemical identity of [$^{64}$Cu]AJ698 by co-elution with its non-radioactive counterpart [$^{nat}$Cu]AJ698. The identical retention times and peak profiles confirm successful and specific incorporation of $^{64}$Cu into the AJ698 peptide.

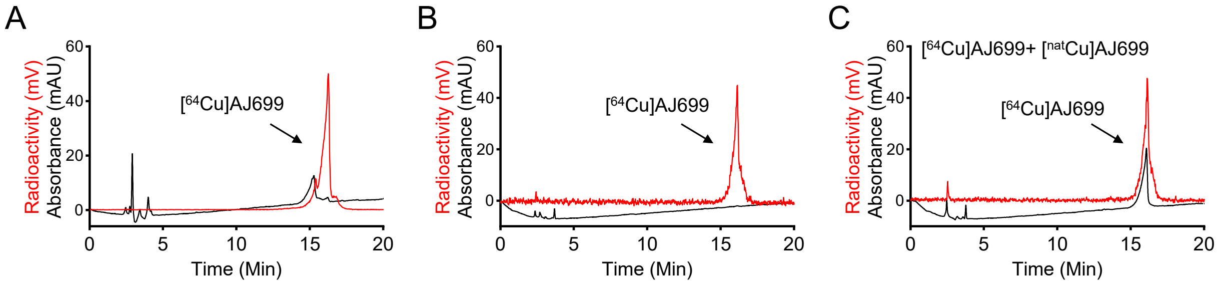


**Figure S20. Radiolabeling and quality control of [$^{64}$Cu]AJ699. A)** HPLC chromatogram of radiotracer purification showing a decay-corrected radiochemical yield of >95%, confirming efficient labeling of AJ699. **B)** HPLC chromatogram representing quality control analysis of [$^{64}$Cu]AJ699 prior to further biological evaluation. **C)** HPLC chromatogram confirming the chemical identity of [$^{64}$Cu]AJ699 by co-elution with its non-radioactive counterpart [$^{nat}$Cu]AJ699. The identical retention times and peak profiles confirm successful and specific incorporation of $^{64}$Cu into the AJ699 peptide.

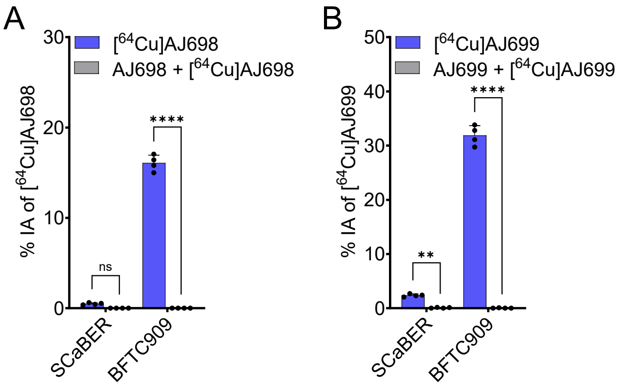


**Figure S21. In vitro evaluation of [$^{64}$Cu]-labeled peptides in bladder cancer cells. A)** [$^{64}$Cu]AJ698, **B)** [$^{64}$Cu]AJ699 binding (%IA) in bladder cancer cells. Cells were incubated with 0.2 µCi of radiotracer at 4 °C for 1 h. Co-incubation with 1 µM of the corresponding non-radioactive peptide (blocking dose) significantly reduced radiotracer uptake, confirming PD-L1–specific binding.

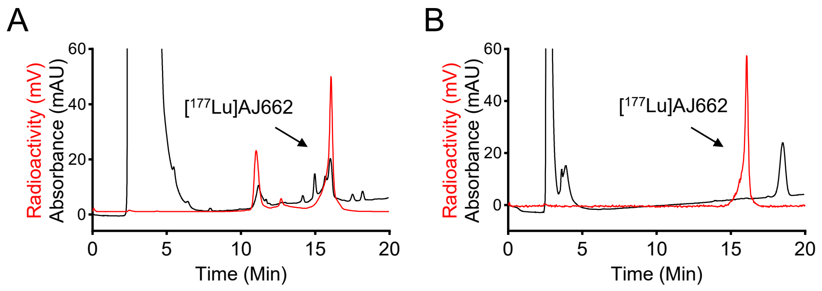


**Figure S22. Radiolabeling and quality control of [$^{177}$Lu]AJ662. A)** HPLC chromatogram of radiotracer purification showing a decay-corrected radiochemical yield of >80%, confirming efficient labeling of AJ662. **B)** HPLC chromatogram representing quality control analysis of [$^{177}$Lu]AJ662 prior to further evaluation.

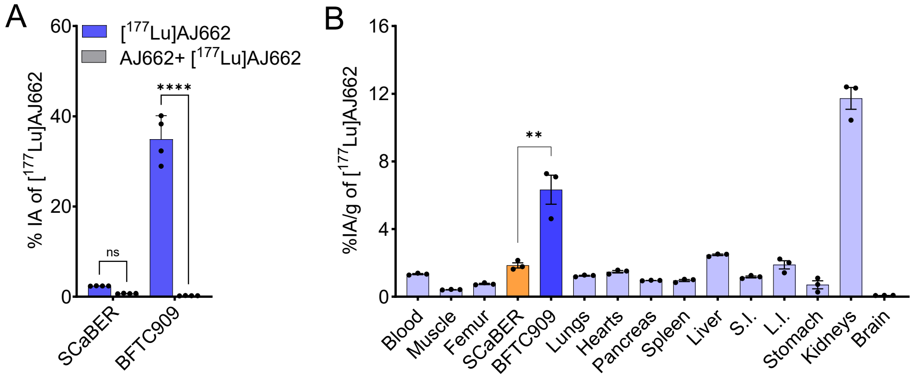


**Figure S23. *In vitro* and in vivo assessment of [$^{177}$Lu]AJ662 in bladder cancer cells and xenograft A)** [$^{177}$Lu]AJ662 binding (percent incubated activity, %IA) to bladder cancer cells. Cells were incubated with 0.5 µCi [$^{177}$Lu]AJ662 at 4°C for 1 hour. [$^{177}$Lu]AJ662 uptake is PDL-1 expression dependent, and co-incubation with 1 µM of non-radioactive AJ662 (blocking dose) significantly reduced radiotracer uptake confirming PDL-1 specificity. **B)** Ex vivo biodistribution analysis in tumor-bearing mice at 120 minutes post-injection of [$^{177}$Lu]AJ662, where S.I. is small intestine and L.I. is large intestine. Data in panels **A** is presented as mean±SD (n=4) and in panels **B** is presented as mean±SEM (n=3). Statistical significance was calculated using two-way ANOVA in panel **A** and unpaired *t* test in **B**; ns, $P \geq 0.05$; **, $P \leq 0.01$; ****, $P \leq 0.0001$.

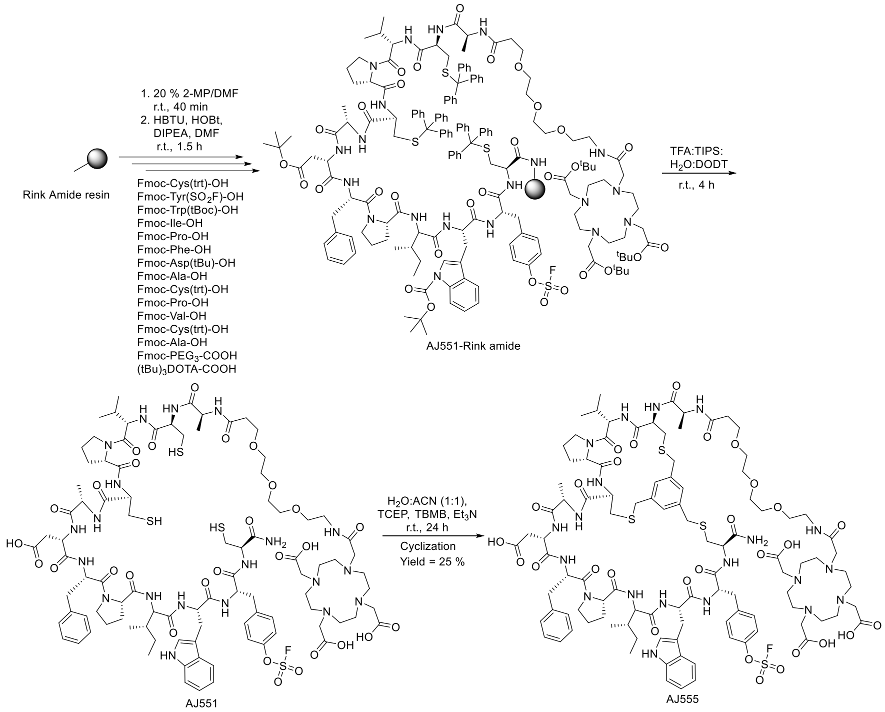


**Scheme 2. Synthesis of Tyr($SO_2F$) containing CD38 binding peptide AJ555.** Automatic solid phase peptide synthesis by adding Fmoc-protected amino acids to Rink amide resin using microwave assisted coupling reaction. The created peptidyl resin was treated with cleavage cocktail to obtain deprotected linear peptide. Liner peptide was cyclized with TBMB in the presence of $Et_3N$ in water:acetonitrile mixture to obtain AJ555.

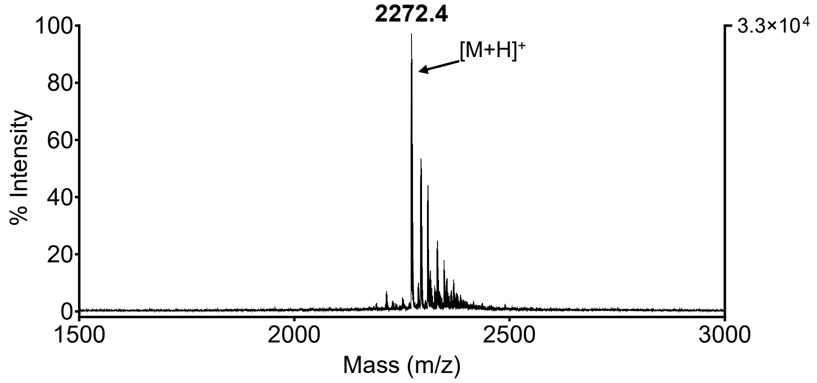


**Figure S24**. Characterization of AJ555 by MALDI-TOF mass spectrometry

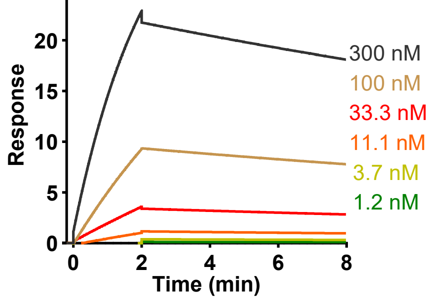


| $K_{on}$ (1/Ms) x$10^4$ | $K_{off}$ (1/s) x$10^{-4}$ | $K_D = K_{off}/K_{on}$ (nM) |
|---|---|---|
| 2.405 | 5.134 | 21.4 |

**Figure S25.** Surface plasmon resonance (SPR) analysis showing nanomolar affinity of AJ555 for recombinant CD38 protein.

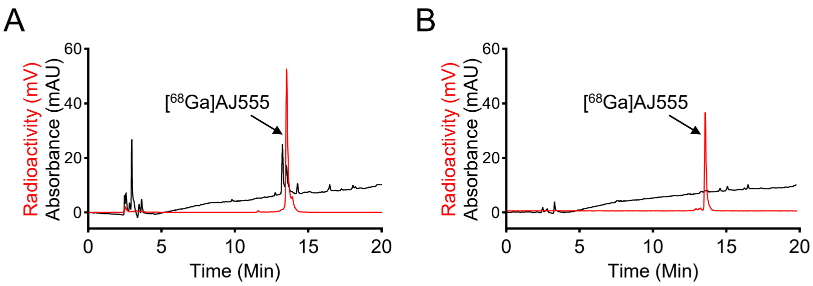


**Figure S26. Radiolabeling and quality control of [$^{68}$Ga]AJ555. A)** HPLC chromatogram of radiotracer purification showing a decay-corrected radiochemical yield of >95%, confirming efficient labeling of AJ555. **B)** HPLC chromatogram representing quality control analysis of [$^{68}$Ga]AJ555 prior to further evaluation.

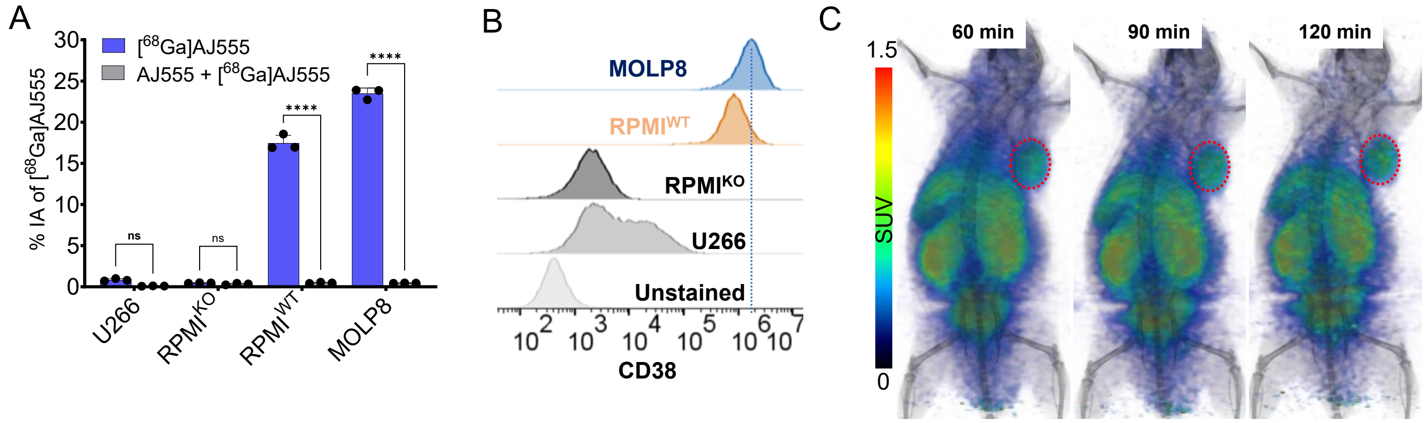


**Figure S27 .** In vitro and in vivo uptake of [$^{68}$Ga]AJ555 **A)** In vitro uptake of [$^{68}$Ga]AJ555 with MM tumor cell lines at 4 °C. Co-incubation with 1 µM of the corresponding non-radioactive peptide (blocking dose) significantly reduced radiotracer uptake, confirming CD38–specific binding. **B)** Flow cytometry assay to evaluate CD38 expression in MM cell lines **C)** In vivo uptake of [$^{68}$Ga]AJ555 in MOLP8 tumor xenograft models using static PET imaging at 60, 90 and 120 min post-tracer injection. Red circle is showing the tumor location. Data in figure **A** is represented as mean±SD (n = 3).

**Table S4**. Ex vivo biodistribution study with MOLP8 tumor bearing NSG mice with [$^{68}$Ga]AJ555 at 2, 60 and 120 min post-tracer injection.

| | Percentage incubated activity per gram (%IA/g) in Mean±SEM (n=5) | | |
|---|---|---|---|
| Tissues | 2 min | 60 min | 120 min |
| blood | 27.82 ± 2.16 | 2.88 ± 0.29 | 1.2 ± 0.21 |
| Femur | 1.3 ± 0.17 | 0.45 ± 0.08 | 0.15 ± 0.02 |
| muscle | 0.5 ± 0.04 | 0.32 ± 0.05 | 0.1 ± 0.03 |
| tumor | 0.41 ± 0.05 | 3.94 ± 0.21 | 2.7 ± 0.29 |
| Lungs | 14.84 ± 1.16 | 2.18 ± 0.27 | 0.95 ± 0.13 |
| Heart | 9.85 ± 1.05 | 1.1 ± 0.13 | 0.44 ± 0.07 |
| Liver | 7.52 ± 0.84 | 6.1 ± 0.27 | 2.48 ± 0.25 |
| S.I. | 1.16 ± 0.25 | 0.82 ± 0.14 | 0.51 ± 0.20 |
| Stomach | 0.38 ± 0.08 | 0.36 ± 0.01 | 0.11 ± 0.01 |
| Spleen | 1.03 ± 0.16 | 0.47 ± 0.02 | 0.23 ± 0.02 |

| | | | |
|---|---|---|---|
| Pancreas | 2.14 ± 0.28 | 0.5 ± 0.06 | 0.22 ± 0.04 |
| Kidney | 5.55 ± 0.47 | 18.07 ± 1.15 | 13.51 ± 1.51 |
| Tumor/Muscle | 0.82 ± 0.12 | 13.57 ± 1.73 | 24.95 ± 3.76 |
| Tumor/Blood | 0.01 ± 0.001 | 1.4 ± 0.11 | 2.38 ± 0.24 |

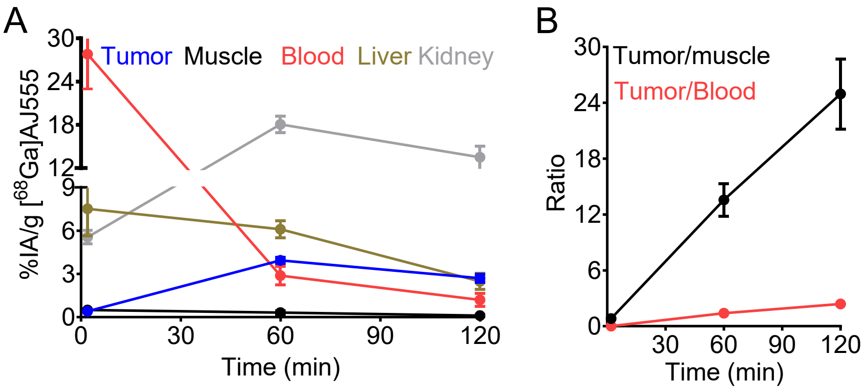


**Figure S28**. **Time activity curve of ex vivo biodistribution study with MOLP8 tumor bearing NSG mice with [$^{68}$Ga]AJ555 A)** %IA/g uptake in tumor, muscle, blood, liver and kidneys and **B)** tumor-to-muscle and tumor-to-blood ratios; data in graphs are represented as mean ± SEM (n = 5)

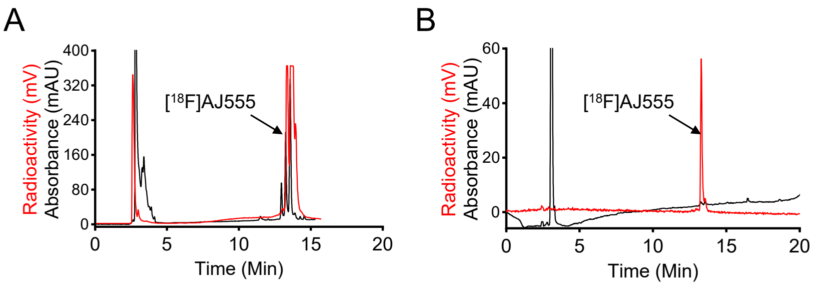


**Figure S29. Radiolabeling and quality control of [$^{18}$F]AJ555. A)** HPLC chromatogram of radiotracer purification showing a decay-corrected radiochemical yield of >95%, confirming efficient labeling of AJ555. **B)** HPLC chromatogram representing quality control analysis of [$^{18}$F]AJ555 prior to further evaluation.

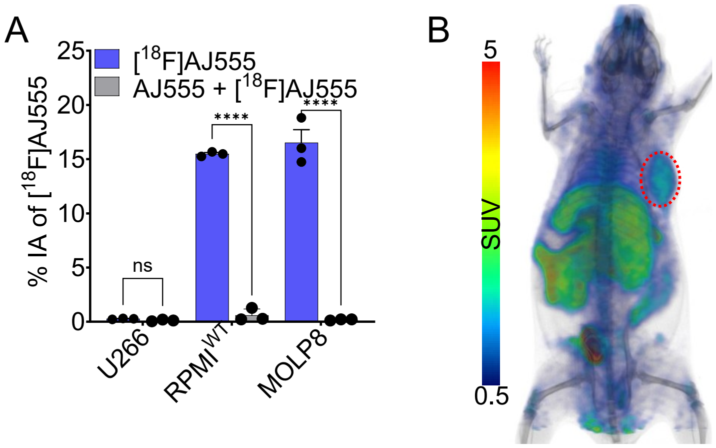


**Figure S30 .** In vitro and in vivo uptake of [$^{18}$F]AJ555 **A)** In vitro uptake of [$^{18}$F]AJ555 with MM tumor cell lines at 4 °C. Co-incubation with 1 μM of the corresponding non-radioactive peptide (blocking dose) significantly reduced radiotracer uptake, confirming CD38–specific binding. **B) I**n vivo uptake of [$^{18}$F]AJ555 in MOLP8 tumor xenograft models using static PET imaging at 60 min post-tracer injection. Red circle is showing the tumor location. Data in figure **A** is represented as mean±SD (n = 3).
